# Temporal shifts in polygenic traits track major epidemics in Western Eurasia

**DOI:** 10.64898/2026.04.07.717059

**Authors:** Flavio De Angelis, Lars Fehren-Schmitz, Carlos Eduardo G. Amorim

## Abstract

Infectious diseases are recognized as one of the strongest selective forces, exerting a profound influence on the genetic makeup of human populations over time. Recently, large-scale genome-wide association studies (GWAS) on immunological traits have underscored the notion that genetic predisposition to infectious diseases likely stems from the contribution of several thousand loci across the human genome. To model the polygenic inheritance of these traits, multiple variants can be combined into polygenic risk scores (PRS), which estimate an individual’s genetic predisposition for a trait. By combining present-day GWAS statistics from large European biobanks with genomic data from more than 3,500 ancient individuals from Western Eurasia, we characterize temporal changes in four highly heritable infectious disease-related traits over a span of 10,000 years. In doing so, we account for variation in these traits across time, space, and genetic ancestries, and demonstrate that the observed patterns cannot be explained by genetic drift alone. Our findings suggest that major disease outbreaks in human history are associated with shifts in polygenic traits in human populations. Specifically, three events – the Justinian Plague, Antonine Plague, and early medieval measles outbreaks – coincide with significant shifts in the polygenic profiles of these traits. Using a Gene Ontology enrichment approach, we show that these shifts involve multiple systemic biological processes, with a consistent emphasis on metabolic pathways modulating immunological responses both directly and indirectly.

## Introduction

The role of infectious diseases (ID) in shaping the evolutionary trajectory of our species has been extensively explored ^1–6^, particularly during periods when infections were the leading cause of mortality and no effective medical treatment existed. Studies in this area recognize pathogen exposure as one of the strongest and most widespread selective forces faced by humans over time ^6,7^. Pathogen-driven selection has shaped genetic diversity in human populations by promoting the persistence of beneficial mutations and the removal of deleterious ones ^5,6,8^. Consistent with this, the human immune system has been subject to pervasive pathogen-imposed selective pressures throughout our evolutionary history ^9–12^. These pressures have contributed to the tuning of multiple immunological pathways, highlighting their central role in host defense and underscoring the importance of human genetic variation in response to infections ^13^.

In spite of the significance of immunological traits in human evolution, genome-wide variation associated with the liability to infection was poorly defined until recently. Only a small percentage of available genome-wide association studies (GWAS) relates to the broad area of infectious disease ^14,15^, and most of the known associations have often been identified in small candidate gene studies focusing on the HLA region ^16^ or other established immune-related loci with large-magnitude effects ^13^. While studies based on candidate loci focused on rare, highly penetrant mutations with large effect sizes, other studies sought to identify genome-wide common variants underlying ID-related traits by considering a polygenic model typical of complex and multifactorial traits ^8,17,18^. As with many other traits, these studies have highlighted the polygenicity of susceptibility to ID in humans ^8,13^. They demonstrate that several thousand loci across the genome – each with typically tiny effects – can effectively contribute to the genetic liability of ID-related complex traits, according to polygenic ^17^ and even more complex modeling approaches ^19^.

The long-lasting interactions between pathogens and their human hosts provide an opportunity to explore the genetic and evolutionary processes related to the spread of ID and their effects on the genetic makeup of ancient human populations. In particular, the significant transitions occurring in the last 10,000 years in West Eurasia in terms of subsistence, demographic expansions, and population admixture ^5,6,20^ enhanced the microbial ability to propagate within and across human groups, ultimately resulting in massive disease outbreaks ^21,22^. The mounting availability of ancient human genomic data allows for the exploration of host-pathogen relationships across critical time transects that overlap with major epidemics and for detecting significant changes in the host genome associated with adaptation to the selective pressure imposed by ID ^6,7,21,23–27^.

The present study aims to investigate temporal changes in polygenic susceptibility to infectious diseases across periods when these represented a major burden for human populations. By combining present-day GWAS-derived statistics and ancient genomic data, we sought to explore the evolution of ID-related traits across the archeological record, adjusting for potential confounding factors related to population structure and geographical location. Because ancient DNA data often have low depth of coverage, resulting in an artifactual excess of homozygotes, we used pseudo-haploid calls, where only a single allele is called instead of a genotype at each position ^28,29^. To model polygenic inheritance of ID-related traits, we combined the effects of multiple variants into Polygenic Risk Scores (PRS), estimating each ancient individual’s genetic potential for an ID-related trait ^30^. Unlike approaches based solely on the analysis of single genes, our approach fully leverages the information in the genomes by considering not only variants of large effect but also small-effect, common genetic variants. Furthermore, we explored the biological processes underlying the PRS associations through a Gene Ontology (GO) enrichment analysis to investigate biological pathways that modulate or are influenced by the immunological response inferred from our PRS-based analysis.

## Methods

### Genome-wide Association Data

Two large-scale cohorts were explored to derive the association data for ID-related traits: the UK Biobank (UKB) and the FinnGen. UKB derives from a population-based research resource containing in-depth genetic and health/lifestyle phenotypic information from over 500,000 UK participants 40–69 at enrollment ^31^. FinnGen is a project combining genotype data from Finnish biobanks and digital health record data from Finnish health registries from N = 377,277 individuals ^32^.

Genome-wide association data for ID-related traits were derived from a pan-ancestry genetic analysis of the UK Biobank (Pan-UKB; Pan-UKB team. https://pan.ukbb.broadinstitute.org. 2020). We used ancestry-specific GWAS statistics for the European genetically defined ancestry group (EUR) comprising N = 420,531 individuals. Out of 7,096 traits related to both sexes, 7,033 were characterized in EUR; from these, 56 were categorized as ID-related (34 coded as Trait Type: *phecode*; Category: *Infectious diseases*; and 22 coded as Trait Type: *icd10*; Category: *Chapter I Certain infectious and parasitic diseases*). We selected those traits with at least 1,000 cases, excluding sub-categorized traits, for a total of 13 traits (supplementary table S1, Supplementary Material).

In addition, we also leveraged FinnGen summary stats (DF9 release) to screen the polygenic architecture of ID-related phenotypes. Among 2,272 traits defined in the DF9 manifest, 95 are categorized as related to infectious and parasitic diseases (Category: *I Certain infectious and parasitic diseases*). We restricted our analyses to 36 traits with at least 1,000 cases (supplementary table S2, Supplementary Material). The FinnGen traits were previously harmonized with ICD codes as reported in ^33^.

Both datasets, including only individuals putatively pertaining to European ancestry, were selected to limit the issues related to the portability of GWAS associations matching the *a priori-defined* ancestry of the target ancient DNA dataset ^34–36^. We note, however, that the portability from contemporary GWAS to ancient samples remains challenging ^37,38^.

### SNP-based heritability and genetic correlation

In order to deal with phenotypes with a reliably heritable basis, we checked their SNP-based heritability, as the proportion of phenotypic variance explained by genome-wide single nucleotide variants (SNVs) ^39^. Furthermore, we determined the genetic correlations across the phenotype panel as the proportion of the heritability that is shared between two traits ^40^. SNP-based heritability and genetic correlation for the ID-related phenotypes were estimated using the Linkage Disequilibrium Score Regression (LDSC) method ^41,42^, considering the HapMap 3 reference panel and pre-computed LD scores based on the 1000 Genomes Project ^43^ reference data for individuals of European ancestry.

### Target Dataset

We collected genetic data from published ancient DNA (aDNA) studies (age ranging from 0-50,000 BP), restricting our analyses to individuals from Western Eurasia – excluding present-day Russia – with pseudo-haploid data for almost 1.24 million SNVs ^44,45^. Pairs previously identified and tagged in the AADR repository as genetically related, low coverage samples (depth of coverage of less than 0.001x), and outliers concerning the local population were removed. Due to the heterogeneous coverages across the dataset, we removed the samples sharing less than 600,000 biallelic variants with the 1240k panel, allowing only 25% variant missingness per sample using *bcftools* ^46^. The resulting dataset comprised 3,555 individuals and 668,017 SNVs. We used NCBI-REMAP to convert GRCh37 coordinates to the GRCh38 release in order to leverage the FinnGen summary statistics.

Due to the impossibility of defining the phenotype for each individual concerning the ID features, we used the average dating reported for each individual as a proxy. These dates were based on either direct radiocarbon dates or well-understood archaeological contexts and reported in the curated dataset as the variable “*Date mean in BP in years before 1950 CE [OxCal mu for a direct radiocarbon date, and average of range for a contextual date]*”. Using these data, we explored the association of the ID-trait genetic liability with the time period when these individuals lived, which, in turn, might match the time of known ID outbreaks. In seeking a balance between resolution and sample size, we focused on individuals from 200 BP to 10,000 BP (N = 2,411) for exploring the PRS associations with time, considering the remote threshold as the Neolithic—a critical time for the ID burden for humans ^5–7^, when most of the already recognized epidemic and pandemic events were triggered by social and demographic factors ^47,48^.

First, we partitioned the time interval in 200-year bins, considering the mean value of the *Date* field reported in the available panel from the AADR v54.1 ^44^. A follow-up analysis was performed by clustering together individuals in 400-year bins. Because of the heterogeneous sample distribution across the time transects, with fewer individuals in remote time, we widened the remote classes as 3,400-4,000; 4,000-8,000; and 8,000-10,000 following the rough cultural boundaries characterizing these periods. We primarily used extended 400-year time bins for our analysis due to the uncertainty in dating each individual. Instead of using radiocarbon dating or contextualized date confidence intervals, we relied on the mean estimate of the date for each individual, which makes it difficult to precisely align them with specific events, such as infectious disease outbreaks.

### PRS analysis

PRSice version 2.3.1.c ^49^ was used to compute PRS. SNPs were clumped based on 250kb windows, based on clump-r2 threshold = 0.1 and clump-p threshold = 1. The step size of the threshold was set to 5E-05, and the starting p-value threshold was 5E-08, using an additive model for regression at each threshold.

Before building association models, PRSice2 allows for a “clumping step” to account for linkage disequilibrium (LD) and reduce redundancy in the SNPs used for PRS calculation. We used a sliding window approach (250 kb) to identify SNPs in LD: within each window, the SNP with the smallest p-value is retained as the index SNP. Other SNPs in the window with r^2^ value greater than the specified threshold (> 0.1) are removed.

Once a list of clumped loci is obtained, the “thresholding step” is used to test different p-value thresholds for SNP inclusion in the PRS. PRSice2 creates multiple PRS models by selecting SNPs based on varying p-value thresholds. We considered p-values ≤ 1, with the starting p-value threshold as 5E-08 and a step size of 5E-05. For each threshold, SNPs with p-values below the threshold are included in the PRS calculation. The PRS for each individual is then calculated by summing the effect sizes of the selected SNPs, weighted by their allele. For each p-value threshold, PRSice2 calculates the PRS for every individual in the target dataset. This results in multiple PRS models, each corresponding to a different p-value threshold. PRSice2 then evaluates the relationship between the PRS and the phenotype using regression models. Specifically, in our strategy, we used a Linear Regression model – as we used Time as a quantitative trait – to obtain beta coefficients and p-values for the association between PRS and Time.

At this stage, PRSice2 allows for the inclusion of covariates in the regression models to control for confounding factors. All PRS were first computed and then covaried for biological sex, the first 20 ancestry-related Principal Components, obtained using Plink v.1.9 ^50^ --pca option (Anc), and latitude/longitude coordinates (Geo). In doing so, we accounted for known confounding factors, and the individual PRS for each sample integrated those factors.

PRSice2 evaluates the association results across all p-value thresholds, and the threshold with the highest R^2^ and significant p-value is selected as the optimal model. False discovery rate (FDR) was applied to correct for multiple testing, accounting for the number of phenotypes and PRS tested, and considering the significant threshold as q = 0.01. The significant association of each PRS with the time-related proxy was tested for the independence of the genetic correlation across the phenotypes, adding the PRS for the correlated traits (if any) as additional covariates to avoid the overfitting of the models due to the genetic correlations across traits. Once the optimal model is identified, the PRS for each individual in our target cohort (the ancient samples) is calculated.

To refine our analysis of PRS associations in relation to pathogen-related burden, we focused on a time range that excludes both the post-antibiotic era, as well as remote prehistoric periods, for which reliable information about the disease landscape is lacking. Accordingly, we removed the present-day individuals (N = 1,116; 31.4%) from the dataset and those whose dating was more than 10,000 years BP (N = 28; 0.8%). This resulted in a sample of 2,411 individuals from 200 to 10,000 years BP. The PRS distributions were normalized to avoid distorting differences in the ranges of values. We performed the Shapiro-Wilk test and the Kruskal-Wallis test on this dataset, seeking to compare the medians of the independent groups represented by individual PRS distribution in each 200- or 400-year time slice. The Shapiro-Wilk test was first performed to assess whether that dataset followed a normal distribution. The null hypothesis (H_0_) is that the data follows a normal distribution, while the alternative hypothesis (H_1_) is that the sample data deviates significantly from a normal distribution. The four fully covaried PRS distributions returned significant results (see Results section); this led us to reject H_0_ and conclude that these PRS distributions are not normally distributed. Accordingly, after median-MAD scaling the PRS distributions to make them less affected by extreme values, we used the non-parametric Kruskal-Wallis test to compare the medians of the independent groups. The null hypothesis (H) states that there is no significant difference in the medians of the groups being compared, while the alternative hypothesis (H) posits that at least one group has a significantly different median.

The Kruskal-Wallis test returned significant results, which prompted us to further investigate the data using a post-hoc Dunn’s test. Dunn’s test is a non-parametric post-hoc test (pairwise comparisons) used after a significant Kruskal-Wallis test to identify which specific groups differ from each other. The test has the following hypotheses: H_0_, the medians of the two groups being compared are equal; and H_1_, the medians of the two groups are not equal. If the adjusted p-value (using Benjamini-Hochberg correction) for a pairwise comparison is less than 0.05, we reject H_0_ and conclude that the medians of the two groups are significantly different. Tests and plots were implemented in R version 4.2.2, using *data.table* (https://cran.r-project.org/web/packages/data.table/data.table.pdf), *dplyr* (https://github.com/tidyverse/dplyr)*, ggplot2* ^51^, and *yarrr* ^52^ packages.

### Assessing the Robustness of PRS-Time Associations

To test for the effect of genetic drift in driving PRS variation, we conducted 10,000 permutations of individual PRS values for each trait by randomly shuffling the effect sizes, effectively disrupting the associations between genetic variation and time. To quantify the shift in PRS, we selected the Spearman correlation coefficient as test statistic, as it does not assume linear relationships. Using the original (non-permuted) data, we calculated the Spearman *rho* to measure the strength of the association between PRS and the time for each trait. We then compared this observed statistic to the distribution of Spearman correlations from the permuted datasets using the *cor* R function with *method = “spearman”,* in order to evaluate how extreme the observed association was under the null hypothesis. If the proportion of permuted datasets where the test statistic exceeds that of the real dataset is large, it suggests the shift we observe in the PRS distributions could be due to drift (i.e., random noise or chance). On the other hand, if the proportion is small, it suggests the shift may be due to non-stochastic processes (i.e., the real trait-genetic association). Specifically, we calculated a p-value as the proportion of permuted datasets with a test statistic as extreme or more extreme than the observed one. A low p-value (e.g., p < 0.05) indicates that fewer than 5% of the permuted datasets replicate the signal and thus that the observed PRS distribution is unlikely to be explained by genetic drift alone.

In addition, we sought to assess whether the observed PRS temporal shifts could be replicated with an independent set of genome-wide variables previously analyzed in the context of a major *Y. pestis* outbreak ^10^. To do so, we built fully-covaried PRS models for the four traits of interest, using the loci reported by Klunk et al. (2022), comprising 356 immune-related genes and 496 loci associated with immune disorders via GWAS. For a complete list of loci, see Klunk et al. (2022).

Finally, to control the potential bias related to the heterogeneous genetic components derived from major socio-demographic shifts throughout the Western Eurasian evolutionary trajectory, a supervised ADMIXTURE analysis ^53^ leveraged all the individuals (n = 3,555) and set as a priori parental groups those defined as Hunter-gatherers, Neolithic, or Late Bronze Age in the AADR v.54.1 ^45^.

### GO Enrichment analysis

We applied a GO Enrichment analysis to interpret the functional significance of gene sets derived from PRS analyses. Rather than evaluating genes in isolation, pathway enrichment analysis identifies biological processes that are statistically overrepresented in a given gene set. The GO enrichment analysis thus serves as a powerful tool for translating gene-level associations into broader system-level insights. The SNVs used to generate each significant PRS association were analyzed for pathway enrichment using PRSet implemented in PRSice v. 2.3.1.c. We used the Molecular Signatures Database (MSigDB) to derive gene sets related to gene ontology terms for biological processes (GO_BioPro) and defined gene boundaries using the human gene annotation ^54^ considering the GRCh37 genome assembly for UKB and GRCh38 for FinnGen. To summarize the overlapping GO terms, we submitted the pathway list to REVIGO ^55^, removing redundant items based on Jiang and Conrath semantic distance ^56^, with a similarity degree of 0.5. To do that, we previously assigned the GO_BioPro term to the proper GO identification number through the R package *BiocManager:GO.db* ^57^, filtering out the terms that did not match the GO ID library. The resulting list was ranked according to the resulting q-values.

## Results

### Genome-wide Association Data and Trait Selection

Genome-wide association data for ID-related traits were obtained from two sources. The first is a pan-ancestry genetic analysis of the UK Biobank (UKB; Pan-UKB team https://pan.ukbb.broadinstitute.org. 2020) ^58^, from where we leveraged ancestry-specific GWAS statistics for ID-related traits for the European genetically defined ancestry (EUR) (accessed in July 2023). The second source is FinnGen (DF9 release, accessed on July 2023), from which we leveraged summary statistics to screen the polygenic architecture of ID-related phenotypes. In both cases, we selected traits for which there were at least 1,000 cases, resulting in 13 and 36 traits respectively.

From the 49 suitable ID-related traits across both biobanks, we selected those that were highly heritable for downstream analyses. To do so, we considered the SNP-based heritability (h2) – i.e., the proportion of phenotypic variance explained by genome-wide single nucleotide variation. In the UKB, h2 of the ID-related traits ranged from 0.0007±0.0011 (“*icd10 A04 Other bacterial intestinal infections*”) to 0.0058±0.0011 (“*phecode 41 Bacterial infection NOS*”) (supplementary table S1, Supplementary Material). FinnGen ID-related traits presented h2 ranging from 0.0002±0.0013 (“*Cholera*”) to 0.019±0.0026 (“*Erysipelas*”) (supplementary table S2, Supplementary Material). After selecting traits with a Z-score of 4 or larger, we identified two traits – “*icd10-B96 Other bacterial agents classified to other chapters*” (h2 = 0.0039±0.0011, Z-score = 4) and “*phecode 41 Bacterial infection NOS*” (h2 = 0.0058±0.0011, Z-score = 5) – with significant h2 in the UKB, considering EUR participants. In FinnGen, we identified five traits with Z-score ≥ 4, namely “*Viral Hepatitis*” (h2 = 0.0061±0.0014, Z-score = 4), “*Erysipelas*” (h2 = 0.0190±0.0026, Z-score = 7), “*Diarrhoea and gastrointestinal infectious origin*” (h2 = 0.0117±0.0015, Z-score = 8), “*Intestinal Infections*” (h2 = 0.0121±0.0014, Z-score = 9), and “*Other Bacterial Diseases*” (h2 = 0.0137±0.0015, Z-score = 9).

As the ID-related traits might share at least partial genetic liability, representing a confounder in association analyses, we used Linkage Disequilibrium Score Regression (LDSC) ^41^ to estimate their genetic correlation (rg), i.e., the proportion of the heritability that is shared between two traits. When looking within each biobank, we found multiple pairs of traits to be significantly correlated (supplementary table S3, Supplementary Material). In the UKB, for instance, “*icd10-B96 Other bacterial agents classified to other chapters*” and “*phecode 41 Bacterial infection NOS*” showed a strongly significant positive correlation (rg = 0.98±0.04, p-value << 1E-10). In FinnGen, rg across the traits ranged between low (“*Erysipelas”* and “*Intestinal Infections”,* rg = 0.22±0.07, p-value < 4E-03) to high values (“*Diarrhoea and gastrointestinal infectious origin”* and “*Intestinal Infections*”, rg = 0.96±0.01, p-value << 1E-10).

### Genetic Liability for Infectious Diseases is Associated with Time

Using GWAS data for the seven ID-related traits with suitable heritability, we constructed PRS association models leveraging a large ancient DNA (aDNA) dataset comprising 3,555 individuals from Western Eurasia ^44^. These PRS estimate an individual’s likelihood of developing a disease or trait by aggregating the effects of multiple genetic variants associated with that phenotype ^30^. As a first step in utilizing PRS to infer the genetic liability for ID-related traits in ancient populations, we explored whether variations in PRS might be associated with major historical and ancient ID outbreaks in Western Eurasia. In the absence of direct phenotypic data for ID exposure, we used time, measured as years before present (BP), as a proxy-phenotype. If the association between PRS and elapsed time is significant, and the model is correctly adjusted for confounding factors, then PRS values for individuals from periods following major outbreaks should significantly differ from those of earlier times, reflecting the selective pressures exerted by these events.

The association between PRS and time was assessed with multiple regression models, including different sets of covariates such as genetic ancestry, geography, and sex. We began with Model 1, which includes no covariates. Model 2 incorporated the first 20 Principal Components (PCs) of genetic data, molecular sex (henceforth referred to as “Sex”) reported in the aDNA data repository ^44^, and geographical location (longitude and latitude coordinates, as reported in the data repository). Subsequent models used a backward removal approach to evaluate the influence of putative confounding factors: Model 3, using geographic coordinates (Geo) and the first 20 PCs (Anc), tested the role of Sex; Model 4, including Sex and Anc, examined the effect of Geo; and Model 5 included Sex and Geo to test for the role of Anc (Table 1).

**Table 1:**
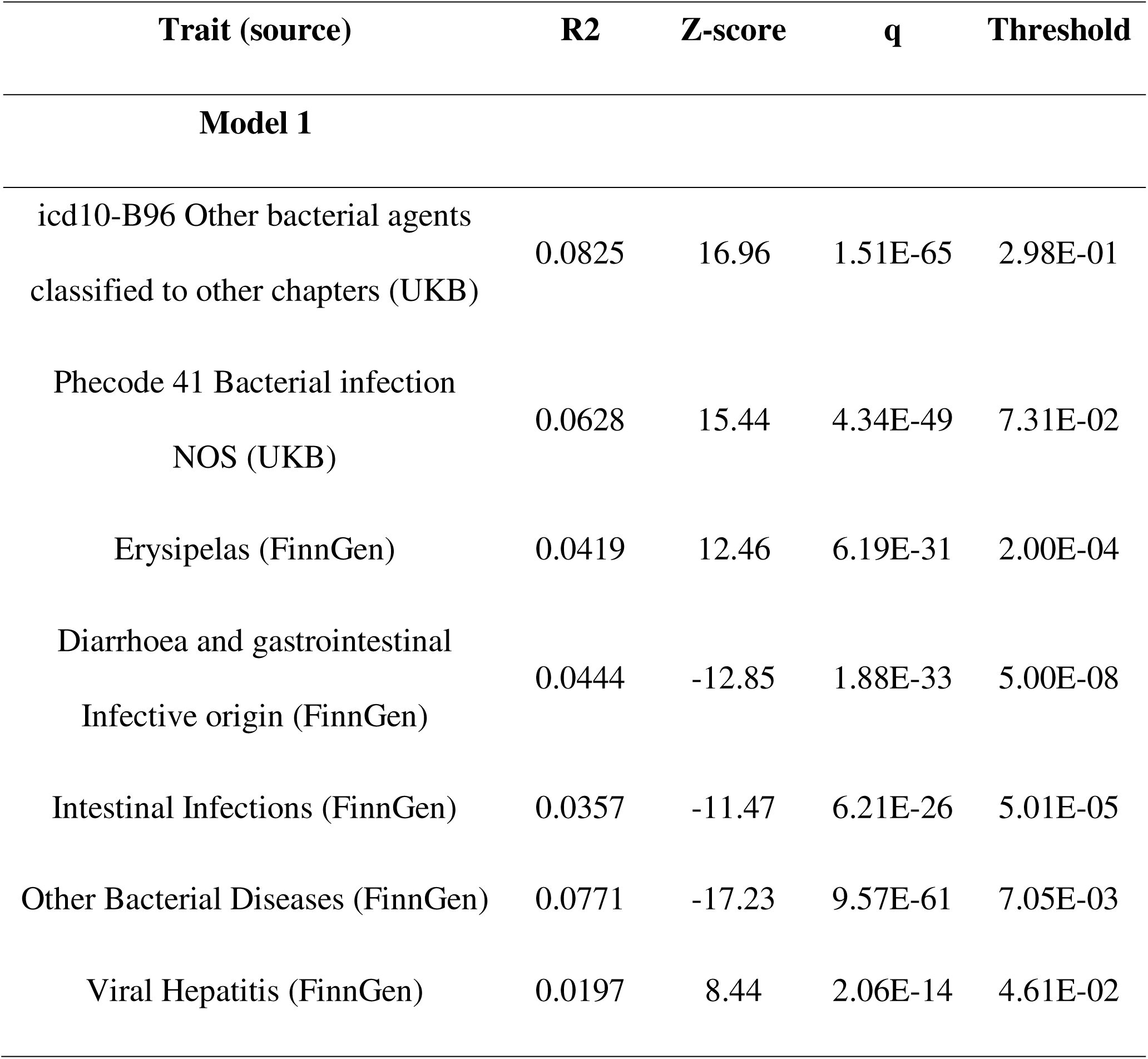
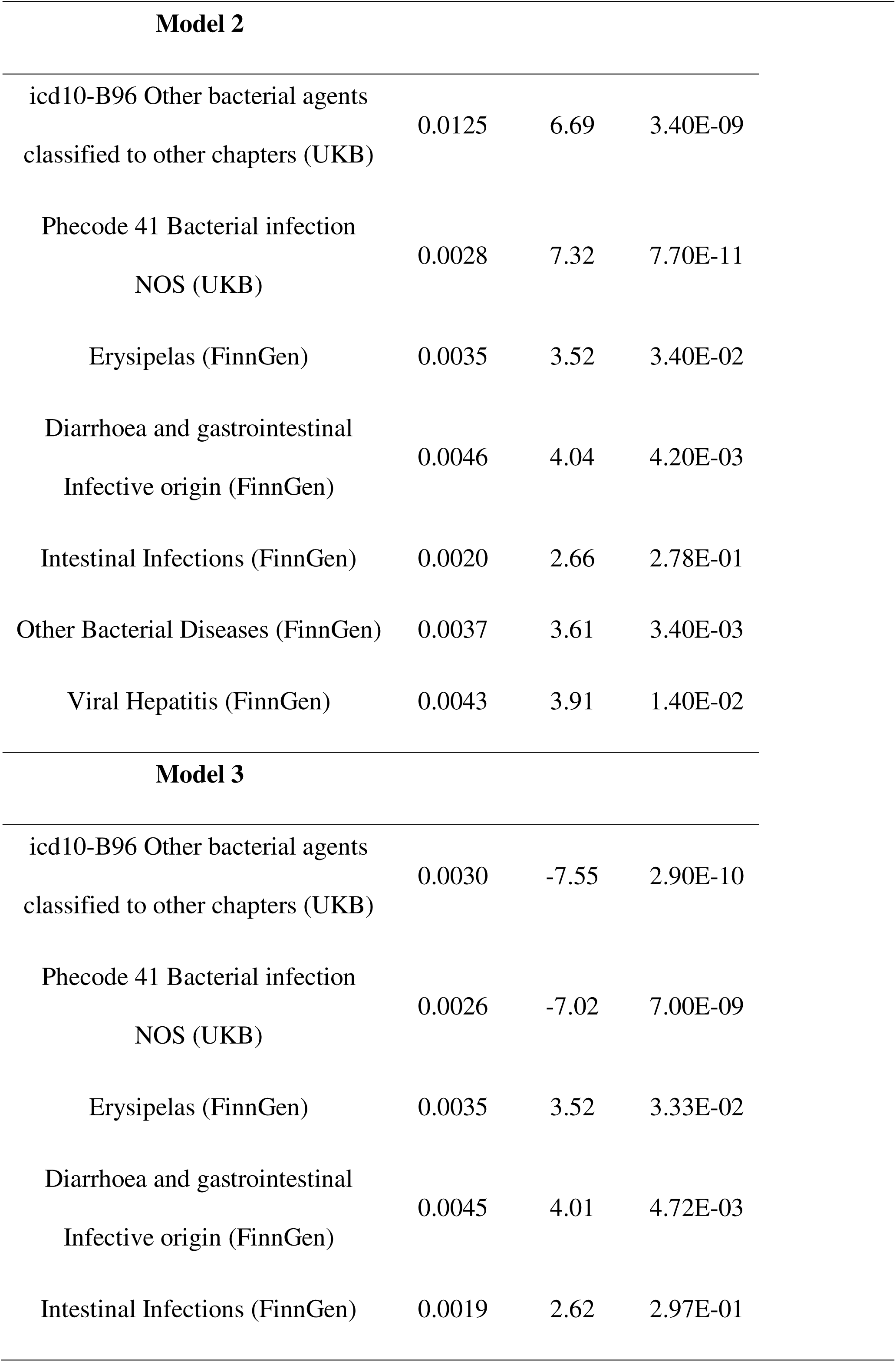
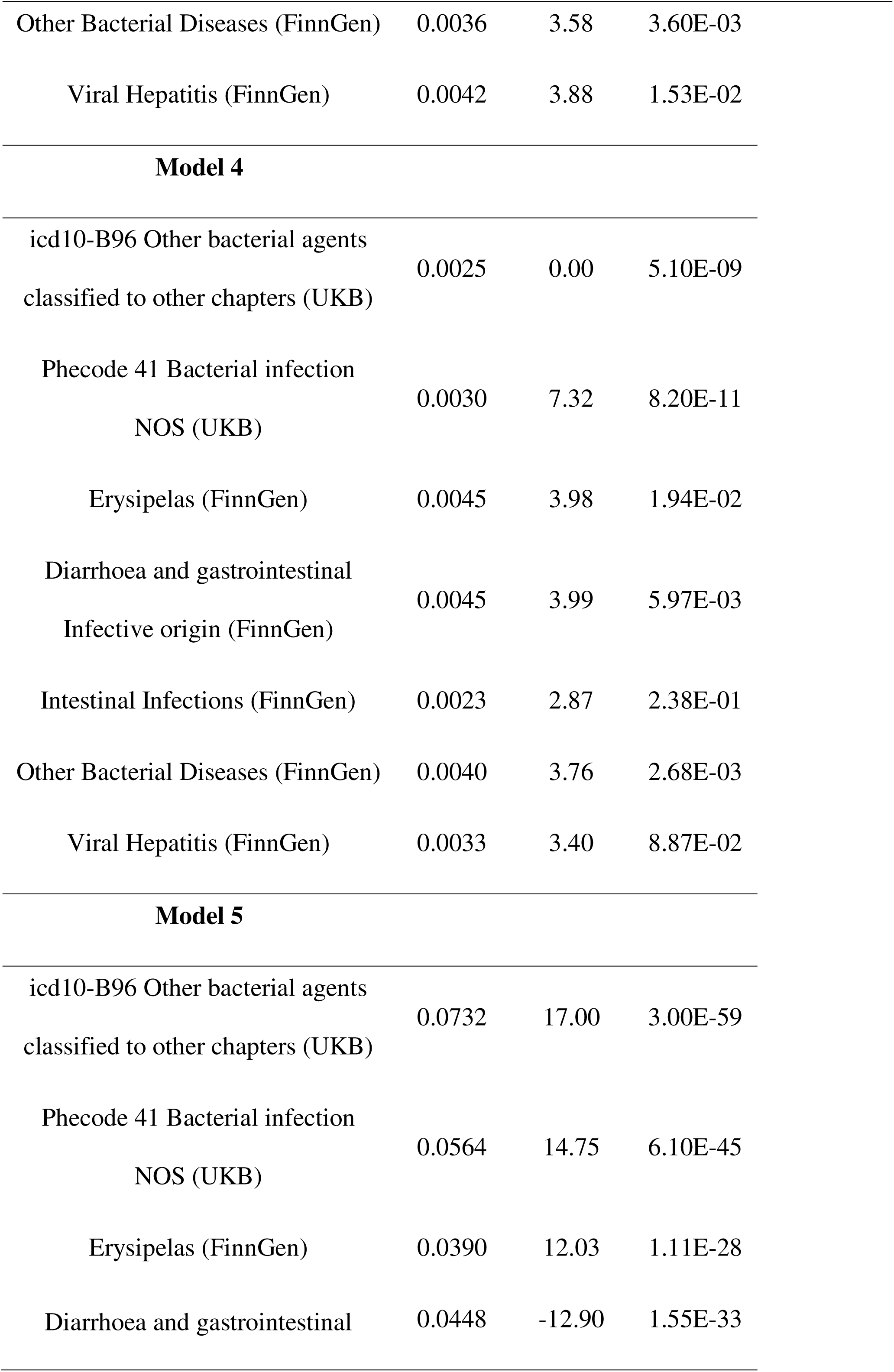
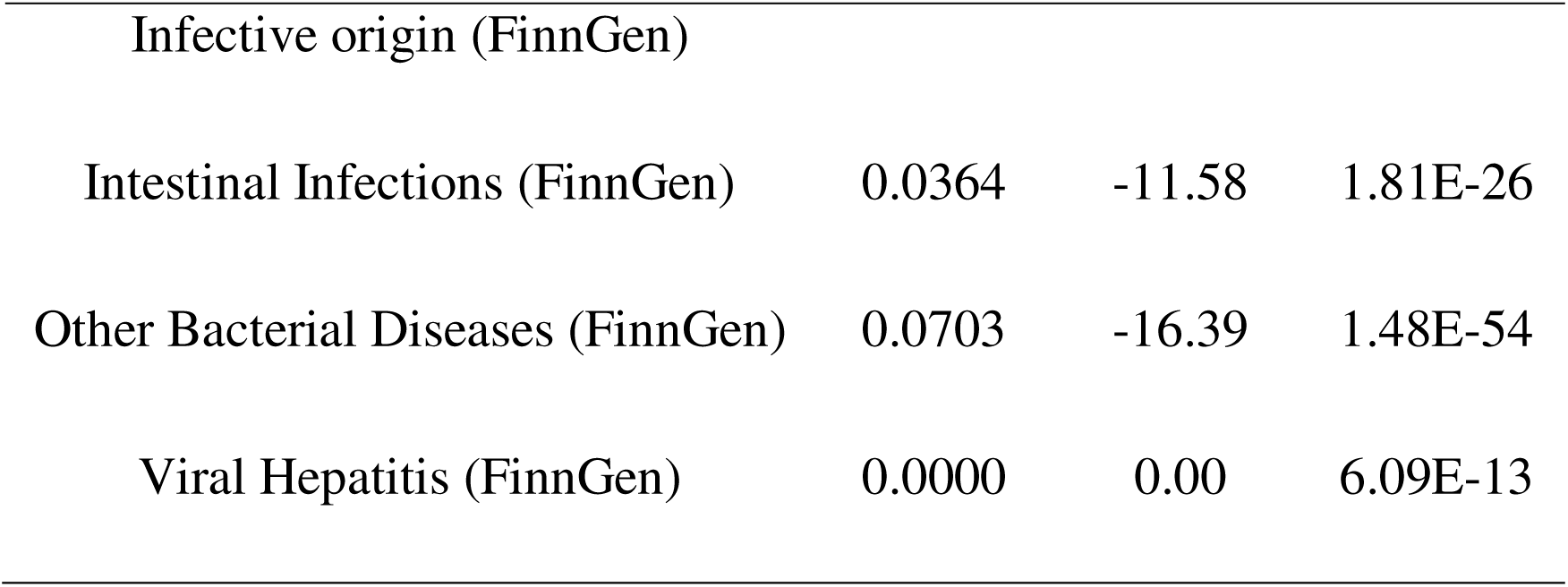
Association models of ID-related trait PRS with time. q represents the expected proportion of false positives according to FDR. R2 = the proportion of phenotypic variance explained by the model. <u>Model 1</u> includes no covariates. The value in the “Threshold” column for Model 1 refers to the optimal p-value threshold for the clumping and thresholding (C+T) approach, which involves testing multiple p-value thresholds to optimize the predictive performance of the resulting PRS. <u>Model 2</u> includes covariates for 20 Principal Components (PCs) of the genetic data, molecular sex, and geographical location (longitude and latitude coordinates). <u>Model 3</u> includes covariates for geographic coordinates and the first 20 PCs. <u>Model4</u> includes covariates for biological sex and the first 20 PCs. <u>Model 5</u> includes covariates for biological sex and geographic coordinates.

We find that PRS for the following four traits are significantly associated with time, even after considering potential confounders: “*icd10-B96 Other bacterial agents classified to other chapters*” and “*phecode 41 Bacterial infection NOS”* from the UKB, and “*Diarrhoea and gastrointestinal infectious origin*” and “*Other Bacterial Diseases*” from FinnGen. Conversely, the trait “*Intestinal Infection*” suffers from a drastic drop in significance when the ancestry components (Anc) are considered in the covariate set (supplementary fig. S1, Supplementary Material), suggesting the role of population stratification in modulating that association. These results indicate that the association models developed for the genetic liability to each of those four traits is consistently significant, demonstrating that the different sets of covariates are not driving the observed associations. Thus, our findings demonstrate that the period in which individuals lived significantly shaped their genetic liability to the ID-related phenotypes.

We note that all the models (to a lesser extent *“Diarrhoea and gastrointestinal infectious origin”*) refer to a non-specific trait, suggesting that the time-specific events driving these associations might have impacted a broad-spectrum genomic architecture responsible for host-pathogen interactions. Thus, this observation makes the evaluation of the genetic correlations across traits compulsory to avoid model overfitting. The model with an additional covariate represented by the PRS for the genetically correlated trait(s) led to a further drop in the q-values of the polygenic models; however, it did not affect their overall significance (Table 2).

**Table 2:** False Discovery Rate for the association models of the four significant ID-related trait PRS with the time proxy-phenotype. Model 1 includes no covariates, while Model 2 incorporates the full set of covariates, including Sex, Ancestry (measured through 20 genetic Principal Components), geographic location (expressed as latitude and longitude), and the PRS for the genetically correlated trait.

| PRS ID trait | Model 1 | Model 2 |
| --- | --- | --- |
| Diarrhoea and gastrointestinal infectious<br>origin | 1.88E-33 | 7.51E-03 |
| Other Bacterial Diseases | 9.57E-06 | 1.59E-03 |
| icd10-B96 Other bacterial agents classified to<br>other chapters | 1.51E-65 | 3.89E-04 |
| Phecode 41 Bacterial infection NOS | 4.34E-49 | 1.25E-08 |

### Evidence for Polygenic Shifts is Associated with Major Infectious Disease Outbreaks

The PRS for the four traits for which the association with time was found (“*icd10-B96 Other bacterial agents classified to other chapters*” and “*phecode 41 Bacterial infection NOS*” from PanUKB; and *“Diarrhoea and gastrointestinal infectious origin*” and “*Other Bacterial Diseases*” from FinnGen) were leveraged to explore their distribution diachronically, considering the known spreads of ID and their outbreaks in Western Eurasia. In doing so, we sought to assess whether the PRS distributions differed across time and whether these shifts coincided with known disease outbreaks. We focused on the 200-10,000 BP timeframe, eliminating individuals living in more remote times due to small sample sizes and those from the post-antibiotic era, therefore reducing the sample size to 2,411 individuals. Model 1, with no covariates, for this reduced target sample is still significant for all four traits (Table 3), whereas the effect of ancestry and the whole covariate set returned only nominally significant (p-value < 0.05) associations (supplementary table S4, Supplementary Material).

**Table 3:**
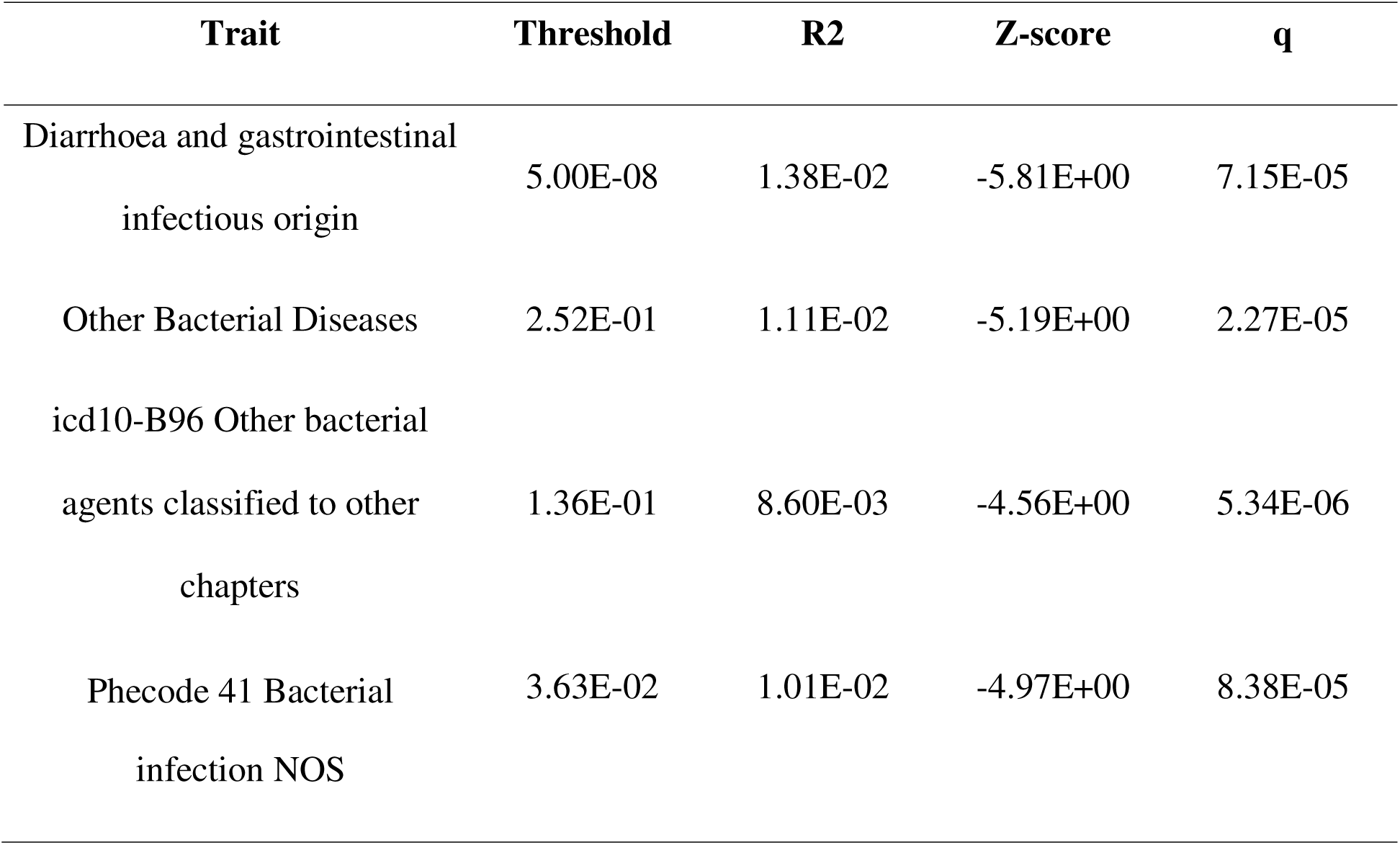
Association Model 1 (without covariates) for four ID-related traits. These models were calculated considering a time range of 200 to 10,000 years before present. R2 represents the proportion of phenotypic variance explained by the model, and q denotes the False Discovery Rate.

To explore the diachronic effect of time (binned in 200- or 400-year intervals) on the PRS distribution, we first applied a Shapiro-Wilk test to evaluate the normality of PRS distributions. Results indicated significant departures from normality across all four traits (“*Diarrhoea and gastrointestinal infectious origin*” W = 0.972, p-value < 0.0001; “*Other Bacterial Diseases*” W = 0.964, p-value < 0.0001; “*Other bacterial agents classified to other chapter*” W = 0.931, p-value < 0.0001; “*Bacterial infection NOS*” W = 0.610, p-value < 0.0001). Consequently, we used a Kruskal-Wallis test to assess whether the PRS distributions significantly differed across time bins. This analysis was performed using the scaled PRS distributions, according to median-MAD scaling, considering the full covaried model (Model 2) with the extended sample (n = 3,555). Individuals were binned according to the date of each sample (radiocarbon date or context-based inference, as reported in the AADR repository). In order to balance the sample sizes across classes, we widened the remote bins beyond 200- and 400-year intervals as following: 3,400-4,000; 4,000-8,000; and 8,000-10,000.

Figure 1 shows the PRS distributions across 400-year bins for the ID-related trait that showed a significant association with time. The PRS distributions for “*Phecode 41Bacterial infection NOS*” differed significantly across bins according to a Kruskal-Wallis test (p-values = 6.84E-04 and 3.58E-05 for the 200-year and 400-year bins respectively). Similarly, PRS distributions for “*icd10-B96 Other bacterial agents classified to other chapters*” also presented significant differences across bins (p-values = 1.71E-03 and 4.86E-05, for the 200- and 400-year bins, respectively). The PRS distributions for the FinnGen-defined traits are consistent with the UKB-based ones. PRS distributions for “*Diarrhoea and gastrointestinal infectious origin*” showed significant differences in the diachronic distribution according to a Kruskal-Wallis test both considering 200-year (p-value = 2.2E-16) and 400-year bins (p-value = 2E-16). Similarly, *“Other Bacterial Diseases*” PRS presented differences across the time transect (p-values = 2.6E-02 and 2.84E-02, for 200- and 400-year bins, respectively).

**Fig. 1:**
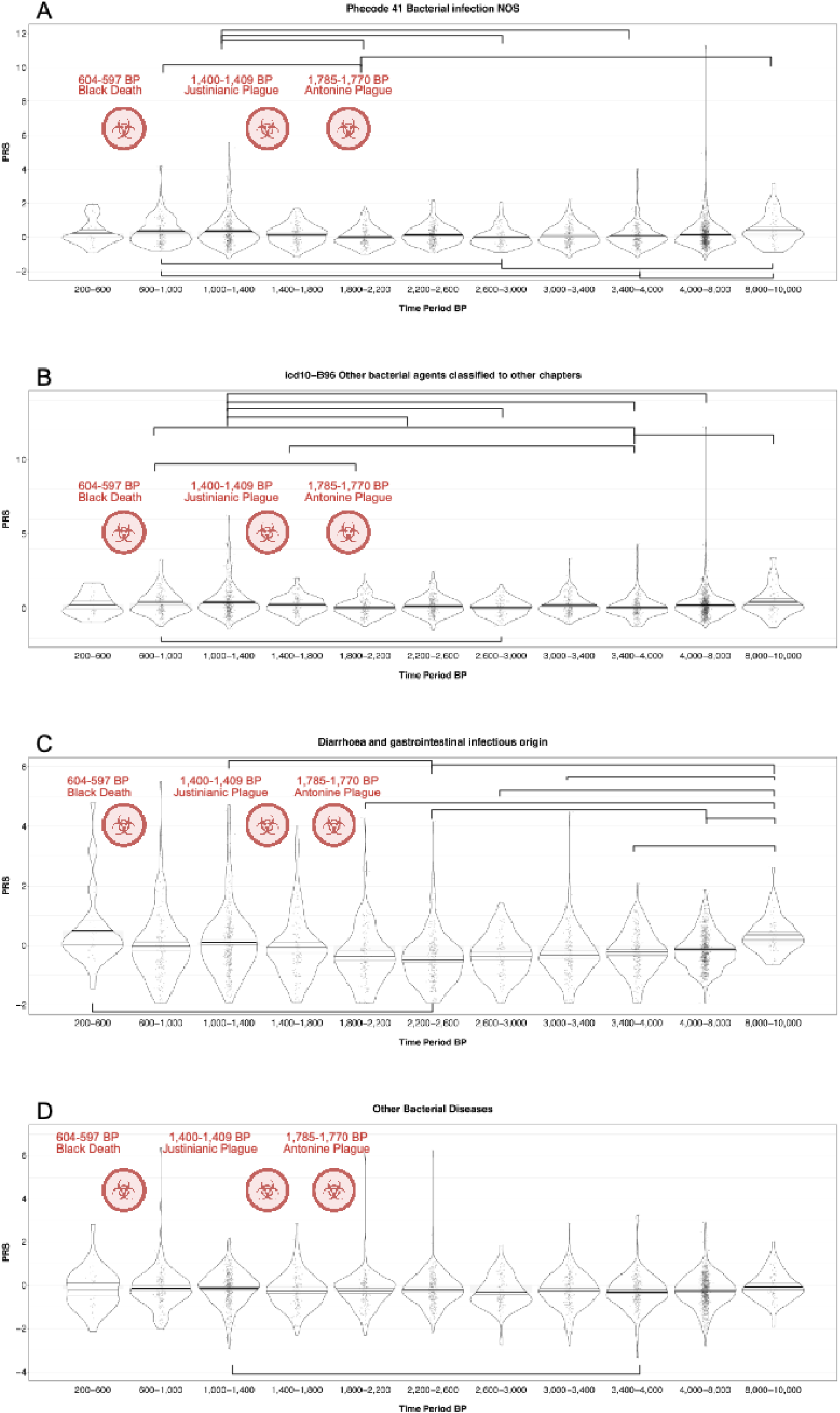
Distribution of polygenic risk scores (PRS) for four infectious disease-related traits across 400-year time intervals. The X-axis represents the 400-year time intervals in years before present, while the Y-axis shows PRS values. A) PRS distribution for the UK Biobank (UKB) trait “*Phecode 41: Bacterial infection, not otherwise specified (NOS)*”. B) PRS distribution for the UKB trait “*ICD-10 B96: Other bacterial agents classified to other chapters*”. C) PRS distribution for the FinnGen trait “*Diarrhoea and gastrointestinal infectious origin*”. Only the nine top-ranking significant associations are represented. D) PRS distribution for the FinnGen trait “*Other Bacterial Diseases*”. The larger biohazard signs refer to the main ID outbreaks discussed in the text or manifestly plague-related outbreaks. Brackets connect Classes that are significantly different according to Dunn’s post-hoc test.

These findings imply that the PRS distribution in at least one time interval was distinct from others. In order to identify the diverging time intervals, we applied a post-hoc test. Dunn’s test highlighted the pairwise differences across the time intervals, accounting for multiple testing (supplementary tables S5-12, Supplementary Material). The evaluation of time transect pairs returning a statistically significant difference showed intriguing findings. Specifically, the 1,400 BP date represented a time threshold that marked significant shifts in PRS distributions for the four traits (supplementary table S13, Supplementary Material). Remarkably, it coincides with the known outbreak of the Justinian Plague that occurred 1,400-1,409 BP, impacting the West Eurasian populations with a death toll estimated at 30-50 million individuals ^59,60^ and, as our results suggest, probably affecting the genetic makeup of the population. Another time point that marks significant shifts in PRS distribution is 1,800 BP (supplementary table S13, Supplementary Material), however only for three of the four traits of interest (“*phecode 41 Bacterial infection NOS”*, “*icd10 B96 Other bacterial agents classified to other chapters”* and “*Diarrhoea and gastrointestinal infectious origin”*). This is consistent with historical sources describing the emergence of a massive epidemic in Europe in 1,770-1,785 BP that suddenly turned into the first known pandemic ravaging the Roman Empire, namely the Antonine Plague – or Plague of Galen, after the description from the Roman Greek physician ^61^. Even the more recent 1,000 BP threshold seemed to be a tipping point for the PRS distributions (supplementary table S13, Supplementary Material). This time point coincides with the outbreak of putative measles/rinderpest etiological agent colonizing and spreading in human populations, causing mass mortality of cattle and people in the early Middle Ages (986–988 CE) in Europe ^62^. Remarkably, the results are overall concordant if we binned the whole sample in 200- or 400-year bins (supplementary tables S5-12, Supplementary Material). We note that the wider 400-year bins better account for dating uncertainty, since we picked the mean of the confidence intervals resulting from the radiocarbon or contextual dating as the best guess for when each individual lived.

The PRSs distributions of each pair of traits – “*icd10 B96 Other bacterial agents classified to other chapters”* vs. *“phecode 41 Bacterial infection NOS*” from UKB; and *“Diarrhoea and gastrointestinal infectious origin”* vs. *“Other Bacterial Diseases”* from FinnGen *–* showed weak but significant positive correlations in the range 200-10,000 BP (Pearson r = 0.0808, p-value < 0.001; and Pearson r = 0.103, p-value < 0.001, respectively). These findings suggest that the putative selective pressure exerted by the disease outbreaks might have modulated the genetic liability of the different traits consistently.

Because genetic drift will cause allele frequencies to fluctuate and the amount of drift accrues with time, we sought to assess whether the association we observe between PRS and time could be due to random drift. To generate null models for the PRS distributions, against which we test whether our observed results can be explained by random drift, we conducted 10,000 permutations of individual PRS values by randomizing the effect sizes, effectively disrupting the genetic variant-trait associations.

To quantify the shift in PRS, we used the Spearman correlation coefficient as test statistic, as it does not assume a linear relationship. We calculated a p-value as the proportion of permuted datasets with a test statistic as extreme or more extreme than the original one with the true effect sizes. We show that the observed PRS distributions for “*phecode 41 Bacterial infection NOS*” (p-value = 0.009) and “*Diarrhoea and gastrointestinal infectious origin*” (p-value = 0.036) are unlikely to be explained by genetic drift alone, whereas for “*icd10-B96 Other bacterial agents classified to other chapters*” (p-value = 0.834) and “*Other Bacterial Diseases*” (p-value = 0.306) that stochastic factors may play a more substantial role (supplementary fig. S2, Supplementary Material).

Having ruled out genetic drift as the sole explanation for the observed PRS shifts, we next investigated whether these temporal PRS shifts could be reproduced using an independent set of loci previously analyzed within the context of the Second Plague Pandemic ^10^. Although some concerns were raised about the study ^63^, we leveraged the full genome-wide SNP set from the Klunk et al. study, which encompasses immune-related variants that are potentially responsive to plague-related pressures. Using this set, we computed fully covariate-adjusted PRS models for the four traits and obtained nominally significant associations for three: “*Diarrhoea and gastrointestinal infectious origin*” (64 loci overlapping with the Klunk et al. SNP set; p-value = 4.79E-15), “*Other Bacterial Diseases*” (60 loci; p-value = 1.6E-03), and “*Phecode 41 Bacterial infection NOS*” (47 loci; p-value = 4.25E-02). The fourth trait, “*ICD10 B96: Other bacterial agents classified to other chapters*” (56 loci), did not yield a significant model (p-value = 8.12E-

01). We then compared PRS distributions between the 1000-1400 BP and 1400-1800 BP time intervals (representing pre- and post-Justinian Plague periods) using the Kolmogorov-Smirnov pairwise test for the three traits with nominally significant models. Despite the limited and uneven variant coverage in the target ancient cohort, we observed significant differences for “*Diarrhoea and gastrointestinal infectious origin”* (D = 0.390; p-value = 2.2E-13) and “*phecode 41 Bacterial infection NOS”* (D = 0.300; p-value = 2.6E-8), thus supporting the trends observed in our models using the extended variant panel. Conversely, *“Other Bacterial Diseases*” (D = 0.069; p-value = 0.726) and “*icd10 B96 Other bacterial agents classified to other chapters”* (D = 0.1; p-value = 0.235) did not show significant differences, likely due to heterogeneous variant coverage between the datasets.

To qualitatively assess potential biases resulting from heterogeneous genetic components that may be unevenly distributed throughout Western Eurasia, we interrogated whether PRS values within each 400-year interval showed systematic PRS gradients structured by geographical location, and whether individuals within each time interval represented a broad range of geographical origins (as opposed to being concentrated in a specific region). We note that our models already account for different covariates, including geographical location. Nonetheless, we sought to provide a qualitative visualization of individual PRS distributions in relation to the analyzed variables. Supplementary figs. S3-6 (Supplementary Material) display the PRS distributions for the four traits of interest, binned into 400-year intervals, with information on geographical location (latitude and longitude) represented by the different colors. If PRS distributions were strongly associated with latitude or longitude, we would expect points to cluster consistently by color. Instead, our results show that the PRS values are scattered across space, with no detectable narrowing of geographical diversity or clustering by PRS values within specific intervals (supplementary figs. S3-6, Supplementary Material). This observation supports the notion that the triggering events impacted the individuals from heterogeneous geographical areas broadly, and that the driver of the observed associations is not geographical location.

Considering a similar approach to what was done with geographical coordinates, we sought to qualitatively assess whether the PRS distribution shifts we detected in the aDNA time series could be explained by broad shifts in genetic ancestry that are known to have occurred in Western Eurasia, such as the Neolithic transition ^64,65^ and the Yamnaya migration ^20,66^. A supervised admixture analysis including individuals associated with Western European Hunter-gatherers, Neolithic, and Late Bronze Age genetic ancestries enabled us to account for genetic components related to human groups experiencing major socio-demographic shifts throughout the Western Eurasian evolutionary trajectory. Our results indicate that no major subsistence strategy changes or massive gene flow from allochthonous peoples moving into Western Eurasia have biased the distributions of the PRS for the four phenotypes, as no significant narrowing of the PRS distribution is evident for specific levels of admixture (supplementary figs. S7-10, Supplementary Material).

### Gene Ontology Enrichment Analysis

To better understand the biological context of the polygenic signals identified in our study, we performed Gene Ontology (GO) enrichment analyses to investigate which functional pathways (GO terms) were overrepresented among the variants included in the PRS models for each trait. While associations with immunity-related processes were expected, given the nature of the traits examined, this analysis allowed us to identify additional biological pathways and processes that may have played a role in the selective events analyzed here, or that could have been indirectly affected by them. This analysis used the full covaried models for the four traits (Model 2). Multiple GO Biological Processes (GO_BioPro) were identified, and their significance was corrected for multiple testing, considering a significant threshold q = 1% (supplementary tables S14-17, Supplementary Material). The redundant GO terms were removed through REVIGO ^55^.

Among the top 10 GO_BioPro related to “*phecode 41 Bacterial infection NOS”* were: *negative regulation of interleukin 5 production* (GO:0032714; P= 8.64E-72; q = 6.69E-68), *positive regulation of interleukin 4 production* (GO: 0032753; P= 1.84E-64; q = 7.13E-61), *interleukin 4 production* (GO: 0032633; P= 1.39E-47; q = 3.59E-44), *ER to Golgi ceramide transport* (GO:0035621; P= 8.42E-40; q = 1.50E-36), *negative regulation of macrophage cytokine production* (GO:0010936; P= 9.71E-40; q = 1.50E-36)*, negative regulation of cytokine production involved in immune response* (GO:0002719, P= 1.26E-38; q = 1.39-35), and *eosinophil migration* (GO:0072677, P= 7.55E-37; q = 7.31E-34).

The GO_BioPro related to “*icd10 B96 Other bacterial agents classified to other chapters”* were enriched for processes related to cellular granulation such as *establishment of pigment granule localization* (GO:0051905; P= 5.47E-52; q = 4.24E-48)*, regulation of Golgi organization* (GO:1903358; P= 1.29E-51; q = 5.00E-48), *pigment granule localization* (GO:0051875; P= 6.38E-51; q = 1.65E-47), and cellular response to hormonal stimuli, such as *cellular response to insulin stimulus* (GO:0032869; P= 7.61E-29; q = 4.16E-26)*, cellular response to peptide hormone stimulus* (GO:0071375; P= 1.36E-24; q = 3.05E-23), and *response to insulin* (GO:0032868; P= 3.51E-23; q = 1.09E-22).

The FinnGen-derived trait associations were also dissected for the GO_BioPro. The associations identified for “*Diarrhoea and gastrointestinal infectious origin”* involved multiple biological processes, including *negative regulation of lipoprotein particle clearance (*GO:0010985; P= 2.64E-63; q= 8.29E-60) and *regulation of Golgi to plasma membrane protein transport* (GO:0042996; P= 2.64E-63; q= 8.29E-60). Significant enrichments related to immune profiles were also identified as *megakaryocyte differentiation* (GO: 0030219; P= 1.17E-15; q = 9.18E-13) and *megakaryocyte development* (GO:0035855; P= 1.17E-15; q = 9.183E-13) and *myeloid cell development (*GO:0061515; P= 1.65E-15; q = 1.15E-12). The loci underpinning the associations related to “*Other Bacterial Diseases”* polygenic risk were enriched for several biological pathways, including *isocitrate metabolic process* (GO:0006102; P= 4.93E-20; q = 3.82E-16); and *flavone metabolic process* (GO:0051552; P= 5.69E-10; q = 6.94E-07).

## Discussion

The interaction between pathogens and host populations is a dynamic process that shaped – and continues to shape – their genomes over time ^5,6,10,67,68^. Infectious diseases have been among the most powerful selective forces in human evolution, leaving a lasting impact on the genetic diversity of our species ^6,22^. Many candidate loci for selection due to pathogen pressure were proposed, mainly comprising immune response genes ^5,6,9,10,13^. However, mounting evidence from large-scale genomic studies, pushed forward by the recent SARS-CoV-2 pandemic, is pointing to an increasingly complex architecture of ID-related traits, featuring significant roles for small-effect variants ^8,17,69–72^. The increasing availability of aDNA data from individuals spanning thousands of years and different geographical locations ^45^ is making it possible to explore the role of the genome-wide common variants involved in ID-related traits by tracking their allele frequency and investigating their roles, even correlating them to ‘when’ and ‘where’ selective events occurred ^5,6,22,73^. Leveraging data from two large-scale biobanks, namely the UKB and FinnGen, we identified ID-related traits showing significant SNP-heritability. Remarkably, we show that those traits are consistently and positively correlated, primarily suggesting the putative sharing of genetic determinants.

Our analyses of the evolution of polygenic ID-related traits through time scrutinized more than 3,500 ancient individuals from Western Eurasia. This sample comprised relatively low coverage data, which was treated as pseudo-haploid. Although we are aware that this approach limits the performance of the PRS calculation, it was previously shown that the imputed data outperform the unimputed data only partially ^38^. Additionally, to prevent the mechanistic loss of effect caused by heterogeneous SNP coverage across samples, we generated a target sample that maximized variant sharing across individuals and limited missingness.

Although the target sample includes extensively analyzed individuals, it is impossible to determine their precise health status or whether they suffered from an infectious disease at the time of death. However, the time period in which these individuals lived provides an indirect proxy for exposure to different outbreaks. Accordingly, we explored whether the polygenic risk for ID-related traits was associated with the chronological context of each individual.

Including the ancestry PCs and the geographic coordinates allowed us to control our models for the effect of the intra-ancestry variability and the covariation with the geographic location. Indeed, we observed that all the tested models suffered from the impact of the genetic ancestry of the samples, which caused a massive drop in the significance of each association model when considered. Uncorrected population stratification was previously recognized as one of the deceiving factors in GWAS, leading to biased or hardly unintelligible results ^74^. The four traits whose models survive the ancestry covariation (“*phecode 41 Bacterial infection NOS”*, “*icd10 B96 Other bacterial agents classified to other chapters”* from UKB and “*Diarrhoea and gastrointestinal infectious origin”, “Other Bacterial Diseases”* from FinnGen) showed robust association models, which are unaffected by geography. Three of these phenotypes refer to non-specific bacterial infections (“*phecode 41 Bacterial infection NOS”*, “*icd10 B96 Other bacterial agents classified to other chapters”* from UKB; and *“Other Bacterial Diseases”* from FinnGen), while the fourth recalls “*Diarrhoea and gastrointestinal infectious origin*”, without specification about the responsible agents (bacteria, viruses, and fungi). This observation suggests that the polygenic determinants of these traits implicated in host-pathogen interactions may be part of an overarching process involving multiple pleiotropic loci, which could provide the background upon which specific candidate loci modulate the pathogen-specific immune responses at the cellular or systemic level ^75^. The exploration of the individual PRS distribution for those traits across diachronic series showed noteworthy results suggesting the role of major ID outbreaks in shaping the genetic liability of ID-related traits through time. Specifically, three events seem to cope with the evidence of significant changes in the polygenic profile of the ID-related traits associations, all relating to the outburst of diseases whose etiological factor was identified in microbes. Even though these events might have been operating only for short bursts, the period when they occurred is associated with a substantial death toll and, consequently, we expect to see a significant shift in the polygenic profile of the ID-related phenotypes associations.

The outbreak of the Justinian Plague around 1,400-1,409 BP could be the driver of the genetic changes observed in Late Antiquity (Fig. 1). The outbreak of this bubonic disease was first reported in the Egyptian town of *Pelusium*, and from there, it quickly spread throughout Western Eurasia, reaching Constantinople in the spring of 542 CE ^76^ up to the British Isles in 544 CE ^77^, continuing for around 200 years and spreading to develop into the First Pandemic. Paleogenetic analyses have accounted for the genome of the Gram-negative bacterium *Yersinia pestis* as responsible for the First Pandemic, revealing the existence of different strains and offering insights into the microbial diversity in Europe between the 6^th^ and 8^th^ centuries CE. Those analyses underscore the evolution of the bacterium over time, including the loss of virulence factor genes later in the pandemic ^77–79^. Many sources provide differing estimates for the plague death toll, with most ranging between 25 and 100 million deaths ^80^. Our results suggest that such a toll would have triggered the shift in the polygenic profile of the phenotypes related to infectious diseases for people living in the periods that followed the outbreak.

Our analyses suggest another tipping point for the genetic liability to ID-related traits around 1,800 BP. Another pandemic, the Antonine Plague, ravaged Western Europe at that time. Despite being referenced as a “plague”, there are no convergent opinions about the etiological agent responsible for the disease and even on the death toll it provided. Evidence suggested it featured a mortality rate of at least 25% across the Roman Empire ^61^. Additionally, the Roman historian *Lucius Cassius Dio* (155-235 CE) estimated from 2,000 to 5,000 deaths per day in Rome at the apex of the outbreak. According to Galen, the symptoms included fever, diarrhea, vomiting, thirstiness, swollen throat, and coughing. More specifically, Galen noted that the diarrhea appeared blackish, which suggested gastrointestinal bleeding ^81^. The initial attribution to smallpox was also supported by foul odor on the breath and exanthema, skin eruptions or rash over the body, distinguished by red and black papules or eruptions ^81,82^. Recent re-evaluation suggested measles as responsible for that massive death toll, based on genetic data reassessing the divergence date between *Measles morbillivirus* (MeV) ^83^, an exclusively human pathogen RNA virus, and its closest relative, the currently eradicated *Rinderpest morbilliviru*s (RPV), a cattle pathogen ^84^. Whatever the pandemic disease impacting humans was, the first detailed description distinguishing smallpox from measles was by Rhazes (860-932), a physician at the hospital in current Baghdad in Roman times ^85^. It is worth mentioning that the gastrointestinal issues and a range of symptoms, including diarrhea and vomiting, were staple signs of the described disease ^86,87^. Remarkably, one of the traits whose PRS distribution was divergent before vs. after the 1,800 BP time point is “*Diarrhoea and gastrointestinal infectious origin”*.

Measles could have reliably been responsible for shaping three of four polygenic signals in recent periods. Our results show that the distribution of the polygenic liability to “*Diarrhoea and gastrointestinal infectious origin”, “icd10 B96 Other bacterial agents classified to other chapters”,* and *“phecode 41 Bacterial infection NOS”* drastically changed around 1,000 CE, when several measles epidemics were reported in Europe, then becoming widespread in South and East Asia, India, and China during the Middle Ages ^85^. As with many infectious diseases, the origin and diffusion of multiple measles outbursts seem plausible in the light of demographic changes: populations large enough to support continuous viral transmission, i.e., 250,000-500,000 individuals, could only rarely exist in previous times ^83^.

While the plague and measles present a plausible explanation for the shifts we observe, other ID outbreaks may also have played a role (supplementary fig. S11, Supplementary Material). However, their limited spread and the lack of reliable mortality data constrain our ability to draw more detailed inferences. Epidemics have historically accompanied major shifts in human society, particularly the transition from hunter-gatherer groups to more densely populated agricultural communities, where both the increased concentration of individuals and the movement of people – through migration, trade, or conflict – have acted as key drivers of disease transmission ^88^. Notably, the 8,000-10,000 BP interval, corresponding to the transition from a hunter-gatherer lifestyle to early agriculture in some parts of Western Eurasia, repeatedly emerges in statistically significant pairwise comparisons of PRS distributions (Fig. 1), underscoring its potential role in shaping genetic susceptibility to infectious diseases. This pattern is even more pronounced in the PRS distribution related to “*Diarrhoea and gastrointestinal infectious origin*” (Fig. 1c; supplementary tables S5 and S13, Supplementary Material). Just as the modern shift toward semi-processed and ultra-processed foods in Western diets has been shown to adversely affect gut microbiota ^89^, the ancient transition from hunter-gatherer/pre-agricultural diets to those based on early farming practices likely disrupted the composition of commensal microbes in the human gastrointestinal tract ^90,91^. This disruption may have had lasting effects on gut health and contributed to the emergence of microbiome-associated diseases.

Beyond the three major ID outbreaks previously discussed and the transition to agriculture, it is important to consider the spread of multiple plague-like diseases that have affected human populations since the Bronze Age in light of findings. Several key events identified through archaeological and genetic research align with the significant shifts in PRS distributions that we observe between 3,000 and 5,000 BP (supplementary fig. S11, Supplementary Material). An example is the oldest known cases of *Y. pestis*-related plague, which were recently discovered in Britain, dating back over 4,000 years ^92^. Around the same time, the Hittite Empire in Anatolia experienced a major epidemic during the mid-14^th^ century BCE, described in historical records by King Muršili II as a prolonged and deadly outbreak ^93^. While originally thought to be plague, evidence now suggests that the disease was likely tularemia, which is caused by *Francisella tularensis* and transmitted through the contact with infected animals or their byproducts ^94^. Additionally, plague outbreaks caused by *Y. pestis* are known to have affected various regions across Asia and Europe between approximately 2,800 and 5,000 years ago, exhibiting varied levels of virulence and modes of transmission ^95^.

Other epidemics also ravaged Western Europe in historical times, although their spread often appears to have been limited or insufficiently documented. Notably, the Plague of Athens (430-426 BCE) was a devastating epidemic that struck during the second year of the Peloponnesian War. It progressed in multiple waves, killing a substantial portion of the Athenian population.

This event might relate to the 2200-2600 BP period, which was among those found to significantly differ from others in our post-hoc statistical analysis (supplementary fig. S11, Supplementary Material). DNA analysis from a mass grave in Athens suggests typhoid fever, caused by *Salmonella enterica*, as the likely pathogen ^96^, though this interpretation remains questioned ^97^.

Another major epidemic, the Plague of Cyprian (249-262 CE), struck Carthage in North Africa and is believed to have entered the Roman Empire through Ethiopia and Egypt ^98^. Named after Saint Cyprian, the bishop of Carthage, who described its symptoms and social impact in his treatise *De Mortalitate*, the plague caused an unknown number of deaths, with ancient sources reporting thousands dying daily in 3^rd^ century Rome, likely due to a viral hemorrhagic fever ^99^. Unfortunately, the temporal resolution of our data, using either 200- or 400-year time slices, does not allow to effectively capture the putative effects of this outbreak.

More recently, English sweating sickness (*Sudor Anglicus*) and syphilis (*lues venerea*) were among the major epidemics that afflicted Western Europe approximately 400-500 years ago ^100,101^. Similarly, the Black Death, one of the most catastrophic pandemics in our history ^102,103^, swept through Europe between 1347 and 1351 CE, causing the death of 25-50 million people, or roughly one-third to one-half of the continent’s population. Unfortunately, our data includes very few samples from this time period, which prevents us from reliably comparing individuals from before and after the events. Nevertheless, the distinctive PRS distribution for the four traits within the 200-600 BP bin (Fig. 1) warrants further investigation using a more targeted analytical approach.

Building on the observed shifts in PRS distributions and their potential link to specific disease outbreaks, we sought to gain further insight into the biological mechanisms underlying these pattern findings. Our GO enrichment analysis identified multiple biological processes implicated in the ID-related traits. Notably, many of the top-ranking GO terms are not limited to (or directly associated with) immune-related processes (supplementary tables S14-17, Supplementary Material), highlighting that the biological pathways emerging from our analysis have broad systemic relevance extending beyond immunology alone ^104,105^.

The GO_BioPro related to “*phecode 41 Bacterial infection NOS”* highlighted processes mainly related to the internalization of pathogens, the induction of apoptosis in infected cells, and the release of cytokines ^106^. Specifically, the GO enrichment analysis highlights biological processes related to the regulation of interleukin (IL)-4 and IL-5. IL-5 exerts pleiotropic activities on various target cells including B cells and eosinophils, which are activated in allergy and can function as antigen-presenting cells ^107^. So far, the abnormal production of a type 2 response cytokines, such as IL-4 and IL-5, has been closely linked to the development of allergic disorders. Type 2 immunity serves essential functions in safeguarding the host, such as defending against helminth parasites and promoting wound repair. However, it is noteworthy that Type 2 immune responses are also implicated in the onset of allergic diseases and autoimmunity through the neutrophils activation. While those cells may have a constructive role in initiating a Type 2 immune response, their participation and activation in later stages are generally undesirable ^108^. Indeed, in their role as effector cells, neutrophils play a role in fostering autoimmune diseases by releasing cascades of cytokines and chemokines ^109^ advocating cooperating immunological cells. It is well known that a de-regulated Type 2 immunity plays a vital role in autoimmune diseases ^110^, and links between the host response to pathogen and autoimmunity are becoming increasingly evident ^5,68,111^, underlining the antagonistic properties of these human characteristics.

The GO_BioPro underpinning the association of “*icd10 B96 Other bacterial agents classified to other chapters”* were enriched for processes related to cellular granulation. Pigmentated intracellular granules relate to multiple and not adequately understood processes. Transient intracytoplasmic green granules in neutrophils have been a matter of discussion about their senescence stage and ability to correlate to the outcome of liver diseases ^112^. Specifically, lipofuscin gives the yellow pigment to granules of lipid containing residues of lysosomal digestion, which could turn to blue-green due to multiple organ failure, especially acute hepatic failure and septic shock secondary to septicemia ^113^. Interestingly, the response to hormone stimuli, specifically insulin, is among the top-ranking GO_BioPro for “*icd10 B96 Other bacterial agents classified to other chapters”*. The role of this peptide hormone in infection is well documented due to the association between critical illness and hyperglycemia. Acute infections are often accompanied by tissue insulin resistance: they contribute to the surge of insulin resistance through enhanced secretion of the counter-regulatory hormones cortisol, glucagon, and even growth hormone ^114^. Moreover, since insulin significantly promotes the macrophage production of TNF-α and IL-6 after the stimulation of Lipopolysaccharides (LPS; i.e., large molecules representing bacterial toxins, especially for Gram-negative microbes), it could also play a critical role in leukocyte function against pathogens ^115,116^. Remarkably, a genome-wide scan for natural selection occurring in the last 10,000 years ^22^ identified candidate loci related to insulin secretion and glucose regulation (mapped to *PTPRV*, *ENSA,* and *MAF*) in the Neolithic, when the immune systems of early farmers responded to the new, pathogen-ridden environment. The Neolithic Transition indeed constituted a major change in people’s subsistence and lifestyle, impacting the genetic makeup of human populations ^117–119^. Still, it was also a critical period for the evolution of immune responses to infectious disease ^7^, which established the genetic adaptations in an antagonistic pleiotropy framework ^111^ with inflammatory traits ^5,7^.

Similarly, the leading GO_BioPro for the associations of PRS for “*Diarrhoea and gastrointestinal infectious origin”* highlighted the role of lipid metabolism and the clearance of lipoproteins. It is well known that infections and inflammation induce various alterations in lipid metabolism that may initially dampen inflammation or fight infection. Changes in lipids and lipoproteins that occur during inflammation and infection – e.g., lipoproteins bind endotoxin, lipoteichoic acid, viruses, and other biological agents and prevent their toxic effects; lipoproteins bind and target parasites for destruction; apolipoproteins neutralize viruses; apolipoproteins lyse parasites – are part of the innate immune response and therefore their production and clearance play an essential role in protecting organisms from the detrimental effects of infection and inflammatory stimuli ^120^. A selection signal was previously detected in loci underpinning the lipid metabolism (*FADS1/2*) in historic time ^22^ or earlier ^6^, often related to the transition to a starch-heavy diet. Consistently, our data could potentially highlight a further role of loci involved in metabolic pathways for modulating the host-pathogen relationship through lipid metabolism ^121^.

Other significant GO_BioPro highlighted the differentiation and development of the megakaryocytes. These cells are known to generate platelets but are also actively involved in the interaction with myeloid cells to promote their migration and stimulate the bacterial phagocytosis of macrophages and neutrophils by producing TNFα and IL-6 ^122^.

Finally, the associations of the PRS for “*Other Bacterial Diseases”* were enriched for isocitrate metabolic process, a leading cellular mechanism in lipid synthesis, cellular defense against oxidative stress, oxidative respiration, and oxygen-sensing signal transduction ^123^. Again, the associations were enriched for the flavone metabolism, which refers to the involvement of molecules able to modulate the activities of several enzymes implicated in immune function ^124^, including cyclooxygenase ^125^, lipoxygenase ^126^, and phosphoinositide 3-kinase ^127^, which feature processes for the response to infective pathological status.

Although our findings contribute to advancing the understanding of complex host-pathogen interactions, we acknowledge several limitations, most notably the challenges related to the portability of PRS to aDNA samples ^37,128,129^ GWAS findings are based on present-day populations whose genetic profiles reflect recent environmental conditions, social structures, and selective pressures that may differ substantially from those that shaped ancient human groups. As such, directly applying GWAS-derived statistics to aDNA data is not straightforward. Over time, processes such as natural selection, genetic drift, and gene flow have reshaped allele frequencies, resulting in genetic differences between ancient and modern populations that can affect the predictive power of PRS ^130,131^. In addition, differences in linkage disequilibrium (LD) patterns between ancient and present-day populations can also influence the portability of PRS (Irving-Pease et al. 2021). Because PRS models rely on the correlation between tag SNPs and causal variants, shifts in LD patterns over time may weaken these correlations, reducing the predictive accuracy of PRS and potentially compromising their portability across time. To mitigate these limitations, our study relies on European-based GWAS and ancient genomes from Western Eurasia, which helps minimize population divergence and provides a more context-appropriate framework.

We also recognize that the nature of host-pathogen interactions has changed over time. In particular, the use of antibiotics may have altered the selective pressures historically exerted by infectious diseases and the genetic architecture of ID-related traits. As medical interventions reduced infection-related mortality and morbidity, the evolutionary dynamics that once shaped host genetic adaptations may have weakened or shifted in direction ^132^. To address these issues, we focus on samples from the pre-antibiotic era, allowing us to examine genetic variation in contexts where pathogen-driven selection likely remained a dominant evolutionary force.

Moreover, the nature of the aDNA adds further complexity to the direct application of PRS. Ancient DNA is often highly fragmented and degraded, resulting in low-quality sequence data with lower coverage than that typically available in modern human genomes (Amorim 2023). These factors may result in issues such as the underrepresentation of rare alleles in ancient samples, base-call errors, and inaccuracies in allele frequency estimates. To mitigate these issues, we applied a strict quality control pipeline, aiming to harmonize the dataset and avoid any mechanistic loss of effect for the variant panel (see Methods).

Finally, the lack of information about each individual’s health status or ID-related markers on the skeletal samples required us to consider the time (years before present) as a proxy to evaluate the diachronic association of the genetic liability to the ID traits. To determine the proxy phenotype value for each individual, we considered the mean dating time of the range reported in the AADR repository. Consequently, we could not pinpoint the exact chronology of the samples. To overcome this limitation and avoid overfitting, we considered both 200- and 400-year bins. The ongoing expansion of available ancient genomic data, more refined dating procedures, and more detailed reports of the osteological characteristics of the analyzed remains may help mitigate these limitations in the future, enabling the application of this approach to larger cohorts sharing comparable infective burden.

In conclusion, our study provides compelling evidence that major events in historical times, particularly those related to infectious diseases, have modulated the polygenic profile of traits associated with infectious disease susceptibility. These findings suggest that the collective influence of numerous common variants has been shaped by shifting pathogen landscapes, environmental pressures, and societal transformations over time. Importantly, this influence appears to operate at a systemic biological level, affecting the complex interplay between the host genome and infectious agents, a dynamic that has evolved throughout history. Such coevolutionary processes likely contributed to shifts in population vulnerability and immunity, reflecting an ongoing arms race between humans and microbes. These insights underscore the importance of considering historical infectious disease challenges when probing the genetic architecture of susceptibility and reveal how systemic and polygenic factors have responded to, and been shaped by, the persistent pressures exerted by infectious diseases over millennia.

## Acknowledgements

We thank Yuval Simons for his input on an early version of this manuscript, Jeremy Berg and Jennifer Blanc for insightful discussions during an earlier phase of the project, and two anonymous reviewers for their constructive feedback and thoughtful suggestions, which helped improve the quality and clarity of this manuscript. Figure 1 and supplementary figure S11 were made using BioRender.com (De Angelis, F. (2025) https://BioRender.com/ebxnqhi; De Angelis, F. (2025) https://BioRender.com/uhmqmca). Research reported in this publication was supported by the National Institute of General Medical Sciences of the National Institutes of Health under award number R35GM142939 for CEGA. FDA was partially supported by the project “MNESYS – a multiscale integrated approach to the study of the nervous system in health and disease”, National Recovery and Resilience Plan, grant number PE0000006, Italian Ministry of University and Research (MUR). The content is solely the responsibility of the authors and does not necessarily represent the official views of the funding institutions.

## Author contributions (CRediT taxonomy)

Conceptualization: F.D.A. and C.E.G.A.; Methodology: F.D.A. and C.E.G.A.; Formal analysis: F.D.A.; Data curation: F.D.A.; Software: F.D.A. and C.E.G.A.; Investigation: F.D.A. and L.F-S; Visualization: F.D.A.; Writing original draft: F.D.A.; Writing – review & editing: All authors; Supervision: C.E.G.A.; Funding acquisition: C.E.G.A.

## Data and code Availability

Ancient genomic data used in this study are available from the Allen Ancient DNA Resource (AADR v54.1). GWAS summary statistics were obtained from the Pan-UK Biobank and FinnGen projects. All processed datasets generated during this study are available from the corresponding author upon reasonable request. Code used for the analyses is available upon request from the corresponding author.

## Additional information

Supplementary Tables: DeAngelis_v2_supptables.xlsx

Supplementary Figures: DeAngelis_v2_supplFigs

## References

1. Daub, J.T., Hofer, T., Cutivet, E., Dupanloup, I., Quintana-Murci, L., Robinson-Rechavi, M., and Excoffier, L. (2013). Evidence for Polygenic Adaptation to Pathogens in the Human Genome. Mol. Biol. Evol. 30, 1544–1558. 10.1093/molbev/mst080.

2. Echaubard, P., Rudge, J.W., and Lefevre, T. (2018). Evolutionary perspectives on human infectious diseases: Challenges, advances, and promises. Evol. Appl. 11, 383–393. 10.1111/eva.12586.

3. Benton, M.L., Abraham, A., LaBella, A.L., Abbot, P., Rokas, A., and Capra, J.A. (2021). The influence of evolutionary history on human health and disease. Nat. Rev. Genet. 22, 269–283. 10.1038/s41576-020-00305-9.

4. King, K.C., Hall, M.D., and Wolinska, J. (2023). Infectious disease ecology and evolution in a changing world. Philos. Trans. R. Soc. B Biol. Sci. 378, 20220002. 10.1098/rstb.2022.0002.

5. Barrie, W., Yang, Y., Irving-Pease, E.K., Attfield, K.E., Scorrano, G., Jensen, L.T., Armen, A.P., Dimopoulos, E.A., Stern, A., Refoyo-Martinez, A., et al. (2024). Elevated genetic risk for multiple sclerosis emerged in steppe pastoralist populations. Nature 625, 321–328. 10.1038/s41586-023-06618-z.

6. Irving-Pease, E.K., Refoyo-Martínez, A., Barrie, W., Ingason, A., Pearson, A., Fischer, A., Sjögren, K.-G., Halgren, A.S., Macleod, R., Demeter, F., et al. (2024). The selection landscape and genetic legacy of ancient Eurasians. Nature 625, 312–320. 10.1038/s41586-023-06705-1.

7. Kerner, G., Neehus, A.-L., Philippot, Q., Bohlen, J., Rinchai, D., Kerrouche, N., Puel, A., Zhang, S.-Y., Boisson-Dupuis, S., Abel, L., et al. (2023). Genetic adaptation to pathogens and increased risk of inflammatory disorders in post-Neolithic Europe. Cell Genomics 3, 100248. 10.1016/j.xgen.2022.100248.

8. Jiang, L., Kerchberger, V.E., Shaffer, C., Dickson, A.L., Ormseth, M.J., Daniel, L.L., Leon, B.G.C., Cox, N.J., Chung, C.P., Wei, W.-Q., et al. (2022). Genome-wide association analyses of common infections in a large practice-based biobank. BMC Genomics 23, 672. 10.1186/s12864-022-08888-9.

9. Brinkworth, J.F. (2017). Infectious Disease and the Diversification of the Human Genome. Hum. Biol. 89, 47–65. 10.13110/humanbiology.89.1.03.

10. Klunk, J., Vilgalys, T.P., Demeure, C.E., Cheng, X., Shiratori, M., Madej, J., Beau, R., Elli, D., Patino, M.I., Redfern, R., et al. (2022). Evolution of immune genes is associated with the Black Death. Nature 611, 312–319. 10.1038/s41586-022-05349-x.

11. Barreiro, L.B., and Quintana-Murci, L. (2010). From evolutionary genetics to human immunology: how selection shapes host defence genes. Nat. Rev. Genet. 11, 17–30. 10.1038/nrg2698.

12. Pankratov, V., Yunusbaeva, M., Ryakhovsky, S., Zarodniuk, M., and Yunusbayev, B. (2022). Prioritizing autoimmunity risk variants for functional analyses by fine-mapping mutations under natural selection. Nat. Commun. 13, 7069. 10.1038/s41467-022-34461-9.

13. Kwok, A.J., Mentzer, A., and Knight, J.C. (2021). Host genetics and infectious disease: new tools, insights and translational opportunities. Nat. Rev. Genet. 22, 137–153. 10.1038/s41576-020-00297-6.

14. Mozzi, A., Pontremoli, C., and Sironi, M. (2018). Genetic susceptibility to infectious diseases: Current status and future perspectives from genome-wide approaches. Infect. Genet. Evol. J. Mol. Epidemiol. Evol. Genet. Infect. Dis. 66, 286–307. 10.1016/j.meegid.2017.09.028.

15. Visscher, P.M., Wray, N.R., Zhang, Q., Sklar, P., McCarthy, M.I., Brown, M.A., and Yang, J. (2017). 10 Years of GWAS Discovery: Biology, Function, and Translation. Am. J. Hum. Genet. 101, 5–22. 10.1016/j.ajhg.2017.06.005.

16. Kulski, J.K., Suzuki, S., and Shiina, T. (2022). Human leukocyte antigen super-locus: nexus of genomic supergenes, SNPs, indels, transcripts, and haplotypes. Hum. Genome Var. 9, 1–15. 10.1038/s41439-022-00226-5.

17. Tian, C., Hromatka, B.S., Kiefer, A.K., Eriksson, N., Noble, S.M., Tung, J.Y., and Hinds, D.A. (2017). Genome-wide association and HLA region fine-mapping studies identify susceptibility loci for multiple common infections. Nat. Commun. 8, 599. 10.1038/s41467-017-00257-5.

18. Butković, A., and Elena, S.F. (2022). Genome-wide association studies of viral infections—A short guide to a successful experimental and statistical analysis. Front. Syst. Biol. 2.

19. Cobey, S. (2020). Modeling infectious disease dynamics. Science 368, 713–714. 10.1126/science.abb5659.

20. Allentoft, M.E., Sikora, M., Refoyo-Martínez, A., Irving-Pease, E.K., Fischer, A., Barrie, W., Ingason, A., Stenderup, J., Sjögren, K.-G., Pearson, A., et al. (2024). Population genomics of post-glacial western Eurasia. Nature 625, 301–311. 10.1038/s41586-023-06865-0.

21. Spyrou, M.A., Bos, K.I., Herbig, A., and Krause, J. (2019). Ancient pathogen genomics as an emerging tool for infectious disease research. Nat. Rev. Genet. 20, 323–340. 10.1038/s41576-019-0119-1.

22. Le, M.K., Smith, O.S., Akbari, A., Harpak, A., Reich, D., and Narasimhan, V.M. (2022). 1,000 ancient genomes uncover 10,000 years of natural selection in Europe. BioRxiv Prepr. Serv. Biol., 2022.08.24.505188. 10.1101/2022.08.24.505188.

23. Marciniak, S., and Perry, G.H. (2017). Harnessing ancient genomes to study the history of human adaptation. Nat. Rev. Genet. 18, 659–674. 10.1038/nrg.2017.65.

24. Gouy, A., and Excoffier, L. (2020). Polygenic Patterns of Adaptive Introgression in Modern Humans Are Mainly Shaped by Response to Pathogens. Mol. Biol. Evol. 37, 1420–1433. 10.1093/molbev/msz306.

25. Immel, A., Key, F.M., Szolek, A., Barquera, R., Robinson, M.K., Harrison, G.F., Palmer, W.H., Spyrou, M.A., Susat, J., Krause-Kyora, B., et al. (2021). Analysis of Genomic DNA from Medieval Plague Victims Suggests Long-Term Effect of Yersinia pestis on Human Immunity Genes. Mol. Biol. Evol. 38, 4059–4076. 10.1093/molbev/msab147.

26. Gao, Z. (2024). Unveiling recent and ongoing adaptive selection in human populations. PLOS Biol. 22, e3002469. 10.1371/journal.pbio.3002469.

27. Hui, R., Scheib, C.L., D’Atanasio, E., Inskip, S.A., Cessford, C., Biagini, S.A., Wohns, A.W., Ali, M.Q.A., Griffith, S.J., Solnik, A., et al. (2024). Genetic history of Cambridgeshire before and after the Black Death. Sci. Adv. 10, eadi5903. 10.1126/sciadv.adi5903.

28. Joseph, T.A., and Pe’er, I. (2019). Inference of Population Structure from Time-Series Genotype Data. Am. J. Hum. Genet. 105, 317–333. 10.1016/j.ajhg.2019.06.002.

29. Amorim, C.E. (2023). Paleogenomics: Ancient DNA in Biological Anthropology. In A Companion to Biological Anthropology (John Wiley & Sons, Ltd), pp. 210–222. 10.1002/9781119828075.ch13.

30. Kullo, I.J., Lewis, C.M., Inouye, M., Martin, A.R., Ripatti, S., and Chatterjee, N. (2022). Polygenic scores in biomedical research. Nat. Rev. Genet. 23, 524–532. 10.1038/s41576-022-00470-z.

31. Bycroft, C., Freeman, C., Petkova, D., Band, G., Elliott, L.T., Sharp, K., Motyer, A., Vukcevic, D., Delaneau, O., O’Connell, J., et al. (2018). The UK Biobank resource with deep phenotyping and genomic data. Nature 562, 203–209. 10.1038/s41586-018-0579-z.

32. Kurki, M.I., Karjalainen, J., Palta, P., Sipilä, T.P., Kristiansson, K., Donner, K.M., Reeve, M.P., Laivuori, H., Aavikko, M., Kaunisto, M.A., et al. (2023). FinnGen provides genetic insights from a well-phenotyped isolated population. Nature 613, 508–518. 10.1038/s41586-022-05473-8.

33. Wu, P., Gifford, A., Meng, X., Li, X., Campbell, H., Varley, T., Zhao, J., Carroll, R., Bastarache, L., Denny, J.C., et al. (2019). Mapping ICD-10 and ICD-10-CM Codes to Phecodes: Workflow Development and Initial Evaluation. JMIR Med. Inform. 7, e14325. 10.2196/14325.

34. Mostafavi, H., Harpak, A., Agarwal, I., Conley, D., Pritchard, J.K., and Przeworski, M. (2020). Variable prediction accuracy of polygenic scores within an ancestry group. eLife 9, e48376. 10.7554/eLife.48376.

35. Matthews, L.J. (2022). Half a century later and we’re back where we started: How the problem of locality turned in to the problem of portability. Stud. Hist. Philos. Sci. 91, 1–9. 10.1016/j.shpsa.2021.10.021.

36. Miao, J., Guo, H., Song, G., Zhao, Z., Hou, L., and Lu, Q. (2023). Quantifying portable genetic effects and improving cross-ancestry genetic prediction with GWAS summary statistics. Nat. Commun. 14, 832. 10.1038/s41467-023-36544-7.

37. Irving-Pease, E.K., Muktupavela, R., Dannemann, M., and Racimo, F. (2021). Quantitative Human Paleogenetics: What can Ancient DNA Tell us About Complex Trait Evolution? Front. Genet. 12.

38. Cox, S.L., Moots, H.M., Stock, J.T., Shbat, A., Bitarello, B.D., Nicklisch, N., Alt, K.W., Haak, W., Rosenstock, E., Ruff, C.B., et al. (2022). Predicting skeletal stature using ancient DNA. Am. J. Biol. Anthropol. 177, 162–174. 10.1002/ajpa.24426.

39. Zhu, H., and Zhou, X. (2020). Statistical methods for SNP heritability estimation and partition: A review. Comput. Struct. Biotechnol. J. 18, 1557–1568. 10.1016/j.csbj.2020.06.011.

40. van Rheenen, W., Peyrot, W.J., Schork, A.J., Lee, S.H., and Wray, N.R. (2019). Genetic correlations of polygenic disease traits: from theory to practice. Nat. Rev. Genet. 20, 567–581. 10.1038/s41576-019-0137-z.

41. Bulik-Sullivan, B., Finucane, H.K., Anttila, V., Gusev, A., Day, F.R., Loh, P.-R., ReproGen Consortium, Psychiatric Genomics Consortium, Genetic Consortium for Anorexia Nervosa of the Wellcome Trust Case Control Consortium 3, Duncan, L., et al. (2015). An atlas of genetic correlations across human diseases and traits. Nat. Genet. 47, 1236–1241. 10.1038/ng.3406.

42. Finucane, H.K., Bulik-Sullivan, B., Gusev, A., Trynka, G., Reshef, Y., Loh, P.-R., Anttila, V., Xu, H., Zang, C., Farh, K., et al. (2015). Partitioning heritability by functional annotation using genome-wide association summary statistics. Nat. Genet. 47, 1228–1235. 10.1038/ng.3404.

43. Auton, A., Abecasis, G.R., Altshuler, D.M., Durbin, R.M., Abecasis, G.R., Bentley, D.R., Chakravarti, A., Clark, A.G., Donnelly, P., Eichler, E.E., et al. (2015). A global reference for human genetic variation. Nature 526, 68–74. 10.1038/nature15393.

44. Mallick, S., Micco, A., Mah, M., Ringbauer, H., Lazaridis, I., Olalde, I., Patterson, N., and Reich, D. (2023). The Allen Ancient DNA Resource (AADR): A curated compendium of ancient human genomes. BioRxiv Prepr. Serv. Biol., 2023.04.06.535797. 10.1101/2023.04.06.535797.

45. Mallick, S., and Reich, D. (2023). The Allen Ancient DNA Resource (AADR): A curated compendium of ancient human genomes. Version 8 (Harvard Dataverse). 10.7910/DVN/FFIDCW 10.7910/DVN/FFIDCW.

46. Danecek, P., Bonfield, J.K., Liddle, J., Marshall, J., Ohan, V., Pollard, M.O., Whitwham, A., Keane, T., McCarthy, S.A., Davies, R.M., et al. (2021). Twelve years of SAMtools and BCFtools. GigaScience 10, giab008. 10.1093/gigascience/giab008.

47. Dowd, J.B., Andriano, L., Brazel, D.M., Rotondi, V., Block, P., Ding, X., Liu, Y., and Mills, M.C. (2020). Demographic science aids in understanding the spread and fatality rates of COVID-19. Proc. Natl. Acad. Sci. 117, 9696–9698. 10.1073/pnas.2004911117.

48. Spernovasilis, N., Markaki, I., Papadakis, M., Tsioutis, C., and Markaki, L. (2021). Epidemics and pandemics: Is human overpopulation the elephant in the room? Ethics Med. Public Health 19, 100728. 10.1016/j.jemep.2021.100728.

49. Choi, S.W., and O’Reilly, P.F. (2019). PRSice-2: Polygenic Risk Score software for biobank-scale data. GigaScience 8, giz082. 10.1093/gigascience/giz082.

50. Purcell, S., Neale, B., Todd-Brown, K., Thomas, L., Ferreira, M.A.R., Bender, D., Maller, J., Sklar, P., de Bakker, P.I.W., Daly, M.J., et al. (2007). PLINK: A Tool Set for Whole-Genome Association and Population-Based Linkage Analyses. Am. J. Hum. Genet. 81, 559–575.

51. Wickham, H. (2009). ggplot2: Elegant Graphics for Data Analysis (Springer New York) 10.1007/978-0-387-98141-3.

52. Phillips, N.D. (2018). YaRrr! The Pirate’s Guide to R.

53. Alexander, D.H., and Lange, K. (2011). Enhancements to the ADMIXTURE algorithm for individual ancestry estimation. BMC Bioinformatics 12, 246. 10.1186/1471-2105-12-246.

54. Yates, A.D., Achuthan, P., Akanni, W., Allen, J., Allen, J., Alvarez-Jarreta, J., Amode, M.R., Armean, I.M., Azov, A.G., Bennett, R., et al. (2020). Ensembl 2020. Nucleic Acids Res. 48, D682–D688. 10.1093/nar/gkz966.

55. Supek, F., Bošnjak, M., Škunca, N., and Šmuc, T. (2011). REVIGO Summarizes and Visualizes Long Lists of Gene Ontology Terms. PLOS ONE 6, e21800. 10.1371/journal.pone.0021800.

56. Couto, F.M., Silva, M.J., and Coutinho, P.M. (2007). Measuring semantic similarity between Gene Ontology terms. Data Knowl. Eng. 61, 137–152. 10.1016/j.datak.2006.05.003.

57. Carlson, M., Falcon, S., Pages, H., and Li, N. (2019). GO. db: A set of annotation maps describing the entire Gene Ontology. R Package Version 3, 10–18129.

58. Karczewski, K.J., Gupta, R., Kanai, M., Lu, W., Tsuo, K., Wang, Y., Walters, R.K., Turley, P., Callier, S., Baya, N., et al. (2024). Pan-UK Biobank GWAS improves discovery, analysis of genetic architecture, and resolution into ancestry-enriched effects. Preprint at medRxiv, 10.1101/2024.03.13.24303864 10.1101/2024.03.13.24303864.

59. Mordechai, L., Eisenberg, M., Newfield, T.P., Izdebski, A., Kay, J.E., and Poinar, H. (2019). The Justinianic Plague: An inconsequential pandemic? Proc. Natl. Acad. Sci. 116, 25546–25554. 10.1073/pnas.1903797116.

60. Horden, P. (2021). plague of Justinian. In Oxford Classical Dictionary 10.1093/acrefore/9780199381135.013.8566.

61. Flemming, R. (2018). Galen and the Plague. In Galen’s Treatise Περ λυπίας (*De indolentia*) in Context (Brill), pp. 219–244. 10.1163/9789004383302_011.

62. Newfield, T.P. (2015). Human–Bovine Plagues in the Early Middle Ages. J. Interdiscip. Hist. 46, 1–38. 10.1162/JINH_a_00794.

63. Barton, A.R., Santander, C.G., Skoglund, P., Moltke, I., Reich, D., and Mathieson, I. (2025). Insufficient evidence for natural selection associated with the Black Death. Nature 638, E19–E22. 10.1038/s41586-024-08496-5.

64. Davy, T., Ju, D., Mathieson, I., and Skoglund, P. (2023). Hunter-gatherer admixture facilitated natural selection in Neolithic European farmers. Curr. Biol. CB 33, 1365–1371.e3. 10.1016/j.cub.2023.02.049.

65. Posth, C., Yu, H., Ghalichi, A., Rougier, H., Crevecoeur, I., Huang, Y., Ringbauer, H., Rohrlach, A.B., Nägele, K., Villalba-Mouco, V., et al. (2023). Palaeogenomics of Upper Palaeolithic to Neolithic European hunter-gatherers. Nature 615, 117–126. 10.1038/s41586-023-05726-0.

66. Papac, L., Ernée, M., Dobeš, M., Langová, M., Rohrlach, A.B., Aron, F., Neumann, G.U., Spyrou, M.A., Rohland, N., Velemínský, P., et al. (2021). Dynamic changes in genomic and social structures in third millennium BCE central Europe. Sci. Adv. 7, eabi6941. 10.1126/sciadv.abi6941.

67. Fumagalli, M., and Sironi, M. (2014). Human genome variability, natural selection and infectious diseases. Curr. Opin. Immunol. 30, 9–16. 10.1016/j.coi.2014.05.001.

68. Kerner, G., Choin, J., and Quintana-Murci, L. (2023). Ancient DNA as a tool for medical research. Nat. Med. 29, 1048–1051. 10.1038/s41591-023-02244-4.

69. Shorter, J.R., Meijsen, J., Nudel, R., Krebs, M., Gådin, J., Mikkelsen, D.H., Nogueira Avelar e Silva, R., Benros, M.E., Thompson, W.K., Ingason, A., et al. (2022). Infection Polygenic Factors Account for a Small Proportion of the Relationship Between Infections and Mental Disorders. Biol. Psychiatry 92, 283–290. 10.1016/j.biopsych.2022.01.007.

70. Bijma, P., Hulst, A.D., and de Jong, M.C.M. (2022). The quantitative genetics of the prevalence of infectious diseases: hidden genetic variation due to indirect genetic effects dominates heritable variation and response to selection. Genetics 220, iyab141. 10.1093/genetics/iyab141.

71. Wendt, F.R., De Lillo, A., Pathak, G.A., De Angelis, F., COVID-19 Host Genetics Initiative, and Polimanti, R. (2021). Host Genetic Liability for Severe COVID-19 Associates with Alcohol Drinking Behavior and Diabetic Outcomes in Participants of European Descent. Front. Genet. 12.

72. Pathak, G.A., Wendt, F.R., Goswami, A., Koller, D., De Angelis, F., and Polimanti, R. (2021). ACE2 Netlas: In silico Functional Characterization and Drug-Gene Interactions of ACE2 Gene Network to Understand Its Potential Involvement in COVID-19 Susceptibility. Front. Genet. 12, 698033. 10.3389/fgene.2021.698033.

73. Mathieson, I., Lazaridis, I., Rohland, N., Mallick, S., Patterson, N., Roodenberg, S.A., Harney, E., Stewardson, K., Fernandes, D., Novak, M., et al. (2015). Genome-wide patterns of selection in 230 ancient Eurasians. Nature 528, 499–503. 10.1038/nature16152.

74. Barton, N., Hermisson, J., and Nordborg, M. (2019). Why structure matters. eLife 8, e45380. 10.7554/eLife.45380.

75. Chattopadhyay, P.K., Roederer, M., and Bolton, D.L. (2018). A deadly dance: the choreography of host–pathogen interactions, as revealed by single-cell technologies. Nat. Commun. 9, 4638. 10.1038/s41467-018-06214-0.

76. Sarris, P. (2022). Viewpoint New Approaches to the ‘Plague of Justinian’*. Past Present 254, 315–346. 10.1093/pastj/gtab024.

77. Keller, M., Spyrou, M.A., Scheib, C.L., Neumann, G.U., Kröpelin, A., Haas-Gebhard, B., Päffgen, B., Haberstroh, J., Ribera i Lacomba, A., Raynaud, C., et al. (2019). Ancient Yersinia pestis genomes from across Western Europe reveal early diversification during the First Pandemic (541–750). Proc. Natl. Acad. Sci. 116, 12363–12372. 10.1073/pnas.1820447116.

78. Feldman, M., Harbeck, M., Keller, M., Spyrou, M.A., Rott, A., Trautmann, B., Scholz, H.C., Päffgen, B., Peters, J., McCormick, M., et al. (2016). A High-Coverage Yersinia pestis Genome from a Sixth-Century Justinianic Plague Victim. Mol. Biol. Evol. 33, 2911–2923. 10.1093/molbev/msw170.

79. Wagner, D.M., Klunk, J., Harbeck, M., Devault, A., Waglechner, N., Sahl, J.W., Enk, J., Birdsell, D.N., Kuch, M., Lumibao, C., et al. (2014). Yersinia pestis and the plague of Justinian 541-543 AD: a genomic analysis. Lancet Infect. Dis. 14, 319–326. 10.1016/S1473-3099(13)70323-2.

80. White, L.A., and Mordechai, L. (2020). Modeling the Justinianic Plague: Comparing hypothesized transmission routes. PLoS ONE 15, e0231256. 10.1371/journal.pone.0231256.

81. Littman, R.J., and Littman, M.L. (1973). Galen and the Antonine plague. Am. J. Philol. 94, 243–255.

82. Gourevitch, D. (2005). The galenic plague: a breakdown of the imperial pathocoenosis. Pathocoenosis and longue durée. Hist. Philos. Life Sci. 27, 57–69.

83. Düx, A., Lequime, S., Patrono, L.V., Vrancken, B., Boral, S., Gogarten, J.F., Hilbig, A., Horst, D., Merkel, K., Prepoint, B., et al. (2020). Measles virus and rinderpest virus divergence dated to the rise of large cities. Science 368, 1367–1370. 10.1126/science.aba9411.

84. Duxbury, E.M., Day, J.P., Maria Vespasiani, D., Thüringer, Y., Tolosana, I., Smith, S.C., Tagliaferri, L., Kamacioglu, A., Lindsley, I., Love, L., et al. (2019). Host-pathogen coevolution increases genetic variation in susceptibility to infection. eLife 8, e46440. 10.7554/eLife.46440.

85. Berche, P. (2022). History of measles. Presse Médicale 51, 104149. 10.1016/j.lpm.2022.104149.

86. Berhe, H.W., Makinde, O.D., and Theuri, D.M. (2019). Co-dynamics of measles and dysentery diarrhea diseases with optimal control and cost-effectiveness analysis. Appl. Math. Comput. 347, 903–921. 10.1016/j.amc.2018.11.049.

87. Simonsen, K.A., and Snowden, J. (2023). Smallpox. In StatPearls (StatPearls Publishing).

88. Piret, J., and Boivin, G. (2021). Pandemics Throughout History. Front. Microbiol. 11.

89. González Olmo, B.M., Butler, M.J., and Barrientos, R.M. (2021). Evolution of the Human Diet and Its Impact on Gut Microbiota, Immune Responses, and Brain Health. Nutrients 13, 196. 10.3390/nu13010196.

90. Alcock, J., Carroll-Portillo, A., Coffman, C., and Lin, H.C. (2021). Evolution of human diet and microbiome-driven disease. Curr. Opin. Physiol. 23, 100455. 10.1016/j.cophys.2021.06.009.

91. Bragazzi, N.L., Del Rio, D., Mayer, E.A., and Mena, P. (2024). We Are What, When, And How We Eat: The Evolutionary Impact of Dietary Shifts on Physical and Cognitive Development, Health, and Disease. Adv. Nutr. 15, 100280. 10.1016/j.advnut.2024.100280.

92. Swali, P., Schulting, R., Gilardet, A., Kelly, M., Anastasiadou, K., Glocke, I., McCabe, J., Williams, M., Audsley, T., Loe, L., et al. (2023). Yersinia pestis genomes reveal plague in Britain 4000 years ago. Nat. Commun. 14, 2930. 10.1038/s41467-023-38393-w.

93. Cate, H.J.H. ten (1969). Hittite Royal Prayers. Numen 16, 81–98. 10.2307/3269758.

94. Trevisanato, S.I. (2007). The ‘Hittite plague’, an epidemic of tularemia and the first record of biological warfare. Med. Hypotheses 69, 1371–1374. 10.1016/j.mehy.2007.03.012.

95. Rasmussen, S., Allentoft, M.E., Nielsen, K., Orlando, L., Sikora, M., Sjögren, K.-G., Pedersen, A.G., Schubert, M., Van Dam, A., Kapel, C.M.O., et al. (2015). Early Divergent Strains of Yersinia pestis in Eurasia 5,000 Years Ago. Cell 163, 571–582. 10.1016/j.cell.2015.10.009.

96. Papagrigorakis, M.J., Yapijakis, C., Synodinos, P.N., and Baziotopoulou-Valavani, E. (2006). DNA examination of ancient dental pulp incriminates typhoid fever as a probable cause of the Plague of Athens. Int. J. Infect. Dis. IJID Off. Publ. Int. Soc. Infect. Dis. 10, 206–214. 10.1016/j.ijid.2005.09.001.

97. Shapiro, B., Rambaut, A., and Gilbert, M.T.P. (2006). No proof that typhoid caused the Plague of Athens (a reply to Papagrigorakis et al.). Int. J. Infect. Dis. 10, 334–335. 10.1016/j.ijid.2006.02.006.

98. Huebner, S.R. (2021). The “Plague of Cyprian”: A revised view of the origin and spread of a 3rd-c. CE pandemic. J. Roman Archaeol. 34, 151–174. 10.1017/S1047759421000349.

99. Kearns, A.L. (2018). A Plague in a Crisis: Differential Diagnosis of the Cyprian Plague and its Effects on the Roman Empire in the Third Century CE.

100. Heyman, P., Simons, L., and Cochez, C. (2014). Were the English Sweating Sickness and the Picardy Sweat Caused by Hantaviruses? Viruses 6, 151–171. 10.3390/v6010151.

101. Tampa, M., Sarbu, I., Matei, C., Benea, V., and Georgescu, S. (2014). Brief History of Syphilis. J. Med. Life 7, 4–10.

102. Glatter, K.A., and Finkelman, P. (2021). History of the Plague: An Ancient Pandemic for the Age of COVID-19. Am. J. Med. 134, 176–181. 10.1016/j.amjmed.2020.08.019.

103. Spyrou, M.A., Musralina, L., Gnecchi Ruscone, G.A., Kocher, A., Borbone, P.-G., Khartanovich, V.I., Buzhilova, A., Djansugurova, L., Bos, K.I., Kühnert, D., et al. (2022). The source of the Black Death in fourteenth-century central Eurasia. Nature 606, 718–724. 10.1038/s41586-022-04800-3.

104. Chicco, D., and Agapito, G. (2022). Nine quick tips for pathway enrichment analysis. PLoS Comput. Biol. 18, e1010348. 10.1371/journal.pcbi.1010348.

105. Barrio-Hernandez, I., Schwartzentruber, J., Shrivastava, A., del-Toro, N., Gonzalez, A., Zhang, Q., Mountjoy, E., Suveges, D., Ochoa, D., Ghoussaini, M., et al. (2023). Network expansion of genetic associations defines a pleiotropy map of human cell biology. Nat. Genet. 55, 389–398. 10.1038/s41588-023-01327-9.

106. Grassmé, H., and Becker, K.A. (2013). Bacterial infections and ceramide. Handb. Exp. Pharmacol., 305–320. 10.1007/978-3-7091-1511-4_15.

107. Gigon, L., Fettrelet, T., Yousefi, S., Simon, D., and Simon, H.-U. (2023). Eosinophils from A to Z. Allergy 78, 1810–1846. 10.1111/all.15751.

108. Egholm, C., Heeb, L.E.M., Impellizzieri, D., and Boyman, O. (2019). The Regulatory Effects of Interleukin-4 Receptor Signaling on Neutrophils in Type 2 Immune Responses. Front. Immunol. 10.

109. Fu, X., Liu, H., Huang, G., and Dai, S. (2021). The emerging role of neutrophils in autoimmune associated disorders: effector, predictor, and therapeutic targets. MedComm 2, 402–413. 10.1002/mco2.69.

110. Hou, X., Chu, J., Liu, S., Jin, S., Sun, J., Wang, H., Li, H., Liu, W., Chai, C., Zhang, S., et al. (2023). Commentary on Updated Insight into the Role of Th2-Associated Immunity in Systemic Lupus Erythematosus. J. Cell. Immunol. Volume 5, 116–119. 10.33696/immunology.5.176.

111. Byars, S.G., and Voskarides, K. (2020). Antagonistic Pleiotropy in Human Disease. J. Mol. Evol. 88, 12–25. 10.1007/s00239-019-09923-2.

112. Gorup, T., Cohen, A.T., Sybenga, A.B., and Rappaport, E.S. (2017). Significance of green granules in neutrophils and monocytes. Proc. Bayl. Univ. Med. Cent. 31, 94–96. 10.1080/08998280.2017.1391045.

113. Soos, M.P., Heideman, C., Shumway, C., Cho, M., Woolf, A., and Kumar, C. (2019). Blue green neutrophilic inclusion bodies in the critically ill patient. Clin. Case Rep. 7, 1249–1252. 10.1002/ccr3.2196.

114. Affinati, A.H., Wallia, A., and Gianchandani, R.Y. (2021). Severe hyperglycemia and insulin resistance in patients with SARS-CoV-2 infection: a report of two cases. Clin. Diabetes Endocrinol. 7, 8. 10.1186/s40842-021-00121-y.

115. Berbudi, A., Rahmadika, N., Tjahjadi, A.I., and Ruslami, R. (2020). Type 2 Diabetes and its Impact on the Immune System. Curr. Diabetes Rev. 16, 442–449. 10.2174/1573399815666191024085838.

116. Tessaro, F.H.G., Ayala, T.S., Nolasco, E.L., Bella, L.M., and Martins, J.O. (2017). Insulin Influences LPS-Induced TNF-α and IL-6 Release Through Distinct Pathways in Mouse Macrophages from Different Compartments. Cell. Physiol. Biochem. Int. J. Exp. Cell. Physiol. Biochem. Pharmacol. 42, 2093–2104. 10.1159/000479904.

117. Bowles, S., and Choi, J.-K. (2019). The Neolithic Agricultural Revolution and the Origins of Private Property. J. Polit. Econ. 127, 2186–2228. 10.1086/701789.

118. Martin-Merino, M. (2021). The Neolithic Revolution : agriculture, sedentary lifestyle and its consequences. Preprint at Cambridge Open Engage, 10.33774/coe-2021-2589h 10.33774/coe-2021-2589h.

119. Childebayeva, A., Rohrlach, A.B., Barquera, R., Rivollat, M., Aron, F., Szolek, A., Kohlbacher, O., Nicklisch, N., Alt, K.W., Gronenborn, D., et al. (2022). Population Genetics and Signatures of Selection in Early Neolithic European Farmers. Mol. Biol. Evol. 39, msac108. 10.1093/molbev/msac108.

120. Feingold, K.R., and Grunfeld, C. (2000). The Effect of Inflammation and Infection on Lipids and Lipoproteins. In Endotext, K. R. Feingold, B. Anawalt, M. R. Blackman, A. Boyce, G. Chrousos, E. Corpas, W. W. de Herder, K. Dhatariya, K. Dungan, J. Hofland, et al., eds. (MDText.com, Inc.).

121. Reynolds, L.M., Dutta, R., Seeds, M.C., Lake, K.N., Hallmark, B., Mathias, R.A., Howard, T.D., and Chilton, F.H. (2020). FADS genetic and metabolomic analyses identify the Δ5 desaturase (FADS1) step as a critical control point in the formation of biologically important lipids. Sci. Rep. 10, 15873. 10.1038/s41598-020-71948-1.

122. Wang, J., Xie, J., Wang, D., Han, X., Chen, M., Shi, G., Jiang, L., and Zhao, M. (2022). CXCR4high megakaryocytes regulate host-defense immunity against bacterial pathogens. eLife 11, e78662. 10.7554/eLife.78662.

123. Tommasini-Ghelfi, S., Murnan, K., Kouri, F.M., Mahajan, A.S., May, J.L., and Stegh, A.H. (2019). Cancer-associated mutation and beyond: The emerging biology of isocitrate dehydrogenases in human disease. Sci. Adv. 5, eaaw4543. 10.1126/sciadv.aaw4543.

124. Chagas, M. do S.S., Behrens, M.D., Moragas-Tellis, C.J., Penedo, G.X.M., Silva, A.R., and Gonçalves-de-Albuquerque, C.F. (2022). Flavonols and Flavones as Potential anti-Inflammatory, Antioxidant, and Antibacterial Compounds. Oxid. Med. Cell. Longev. 2022, 9966750. 10.1155/2022/9966750.

125. Stables, M.J., Newson, J., Ayoub, S.S., Brown, J., Hyams, C.J., and Gilroy, D.W. (2010). Priming innate immune responses to infection by cyclooxygenase inhibition kills antibiotic susceptible and resistant bacteria. Blood 116, 2950–2959. 10.1182/blood-2010-05-284844.

126. Ayola-Serrano, N.C., Roy, N., Fathah, Z., Anwar, M.M., Singh, B., Ammar, N., Sah, R., Elba, A., Utt, R.S., Pecho-Silva, S., et al. (2021). The role of 5-lipoxygenase in the pathophysiology of COVID-19 and its therapeutic implications. Inflamm. Res. 70, 877–889. 10.1007/s00011-021-01473-y.

127. Fleeman, R. (2023). Repurposing inhibitors of phosphoinositide 3-kinase as adjuvant therapeutics for bacterial infections. Front. Antibiot. 2.

128. Carlson, M.O., Rice, D.P., Berg, J.J., and Steinrücken, M. (2022). Polygenic score accuracy in ancient samples: Quantifying the effects of allelic turnover. PLoS Genet. 18, e1010170. 10.1371/journal.pgen.1010170.

129. Le, M.K., Smith, O.S., Akbari, A., Harpak, A., Reich, D., and Narasimhan, V.M. (2022). 1,000 ancient genomes uncover 10,000 years of natural selection in Europe. bioRxiv, 2022.08.24.505188. 10.1101/2022.08.24.505188.

130. Pandey, D., Harris, M., Garud, N.R., and Narasimhan, V.M. (2024). Leveraging ancient DNA to uncover signals of natural selection in Europe lost due to admixture or drift. Nat. Commun. 15, 9772. 10.1038/s41467-024-53852-8.

131. Simon, A., and Coop, G. (2024). The contribution of gene flow, selection, and genetic drift to five thousand years of human allele frequency change. bioRxiv, 2023.07.11.548607. 10.1101/2023.07.11.548607.

132. Kwon, J.H., and Powderly, W.G. (2021). The post-antibiotic era is here. Science 373, 471–471. 10.1126/science.abl5997.

