## Supplemental Figures for "Temporal shifts in polygenic traits track major epidemics in Western Eurasia"

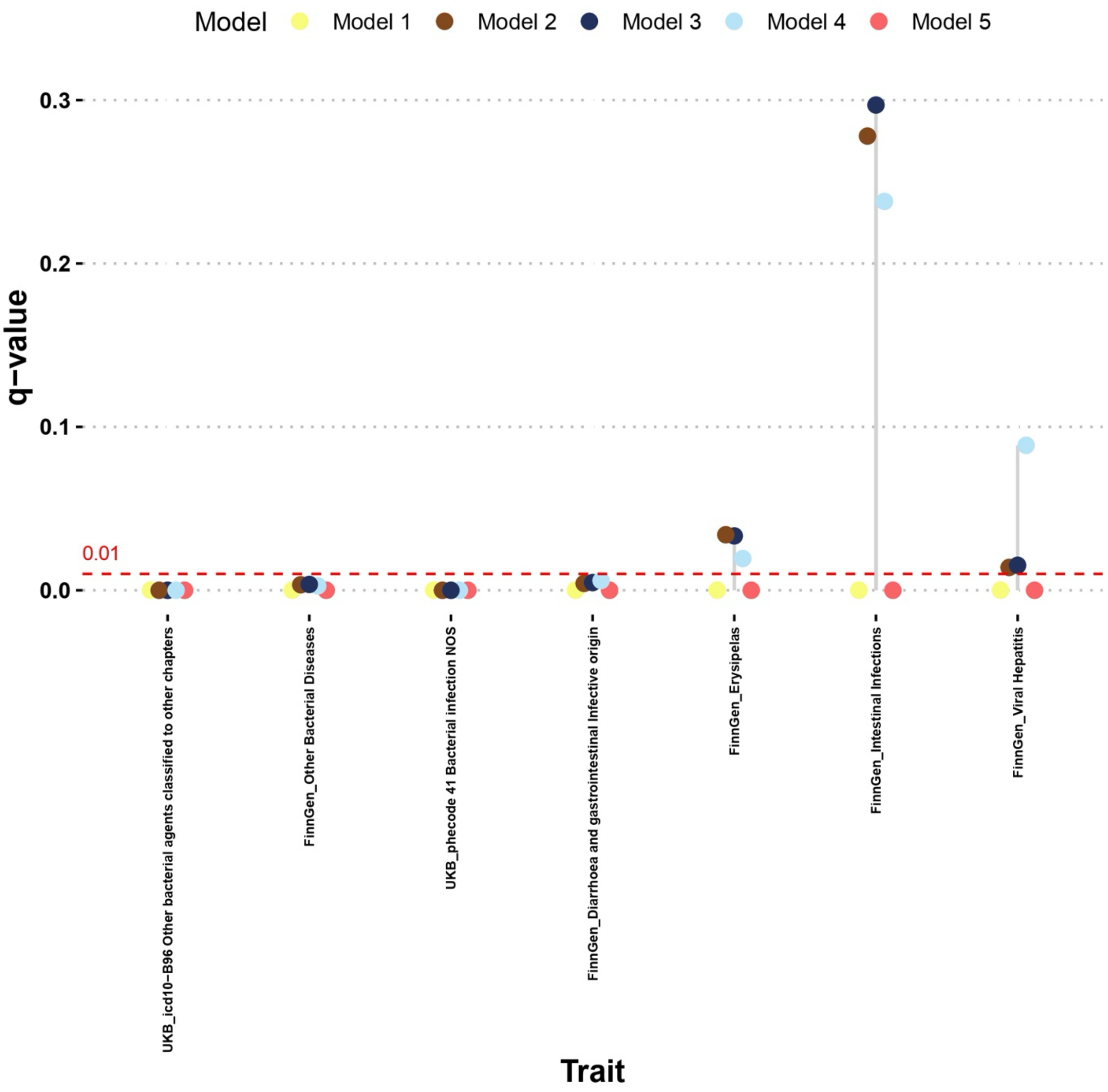


Fig. S1: Cleveland's Dot Plot showing the significance drop across the models. Significance threshold is reported as dashed line (q-value significance threshold = 0.01). Model 1 includes no covariates. Model 2 incorporates the first 20 Principal Components (PCs) of genetic data, molecular sex (Sex), and geographical location (longitude and latitude coordinates). Model 3 leverages geographic coordinates (Geo) and the first 20 PCs (Anc). Model 4 includes Sex and Anc. Model 5 includes Sex and Geo.

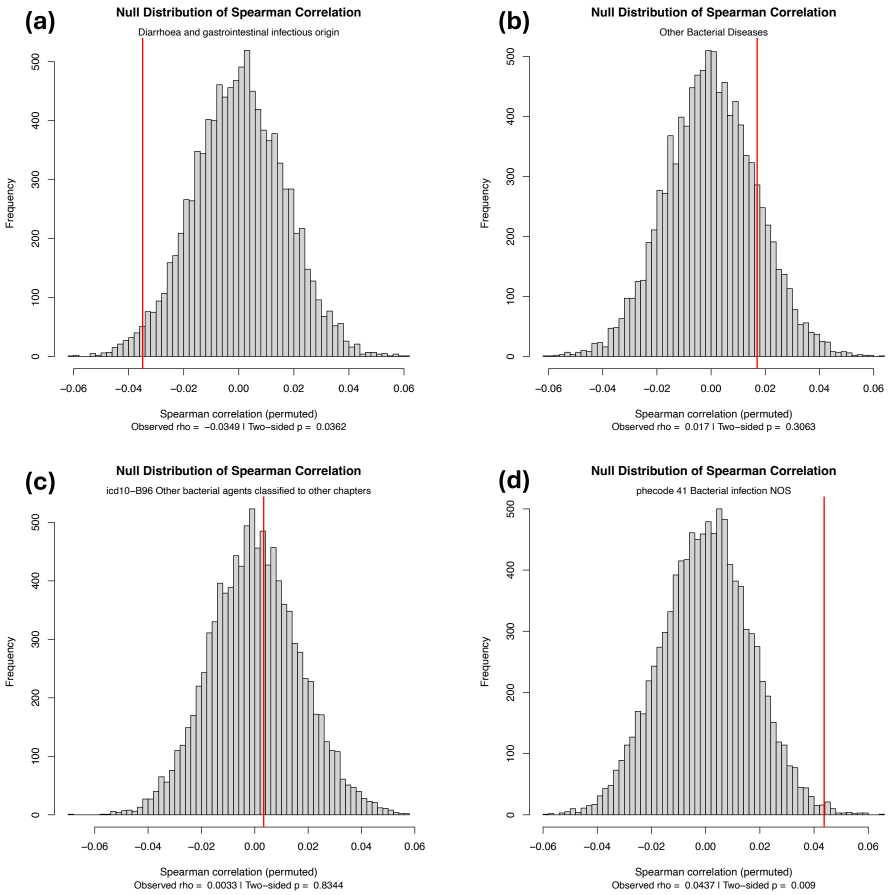


Fig. S2: Comparison of the actual Spearman correlation rho (red line) to the distribution of the statistics from the permuted (n = 10,000) datasets. a) *"Diarrhoea and gastrointestinal infectious origin"*; b) *"Other Bacterial Diseases"*; c) *"icd10-B96 Other bacterial agents classified to other chapters"*; d) *"phecode 41 Bacterial infection NOS"*.

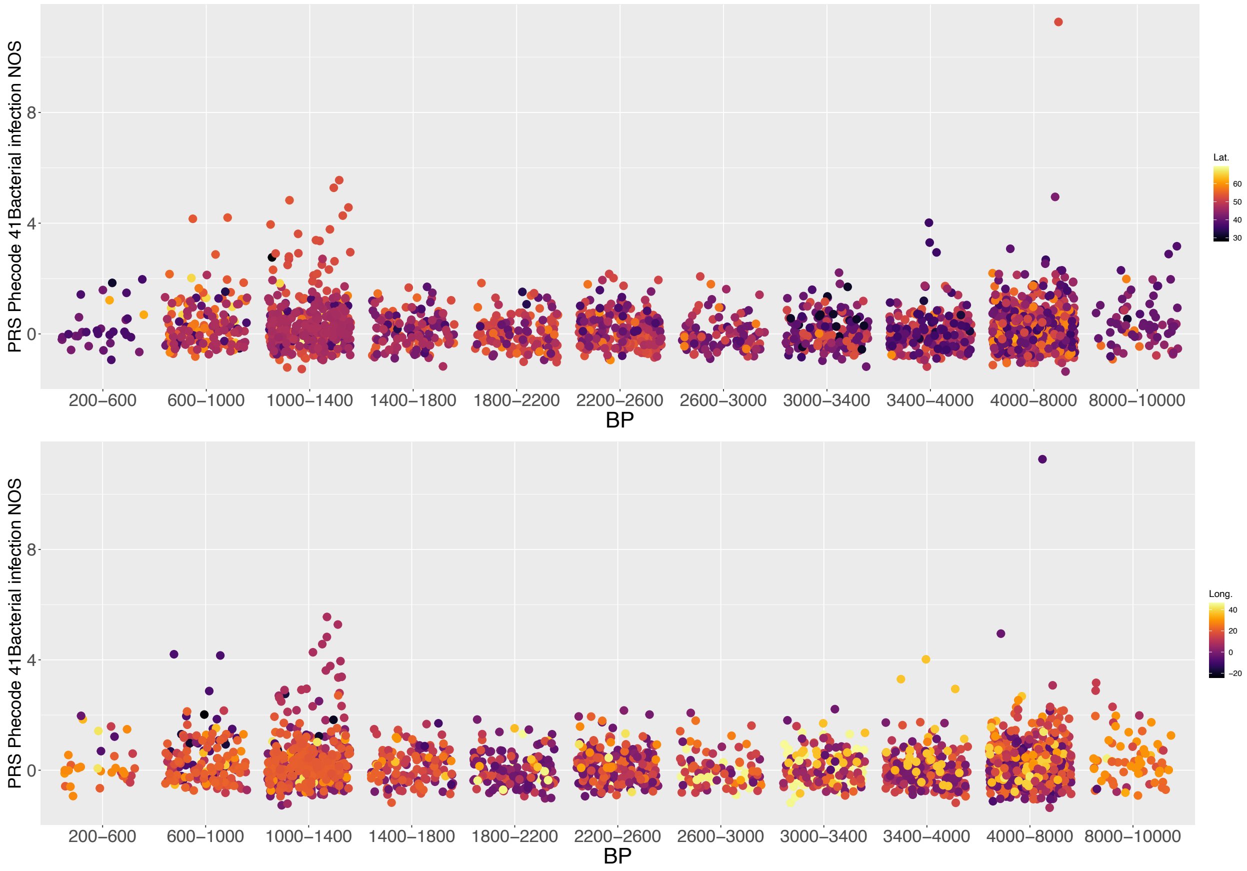


Fig. S3: PRS distribution for *phecode 41 Bacterial infection NOS* binned in 400-year classes. Colors refer to the Latitude (top) and Longitude (bottom) where the individual sample is located according to AADR v.54.1.

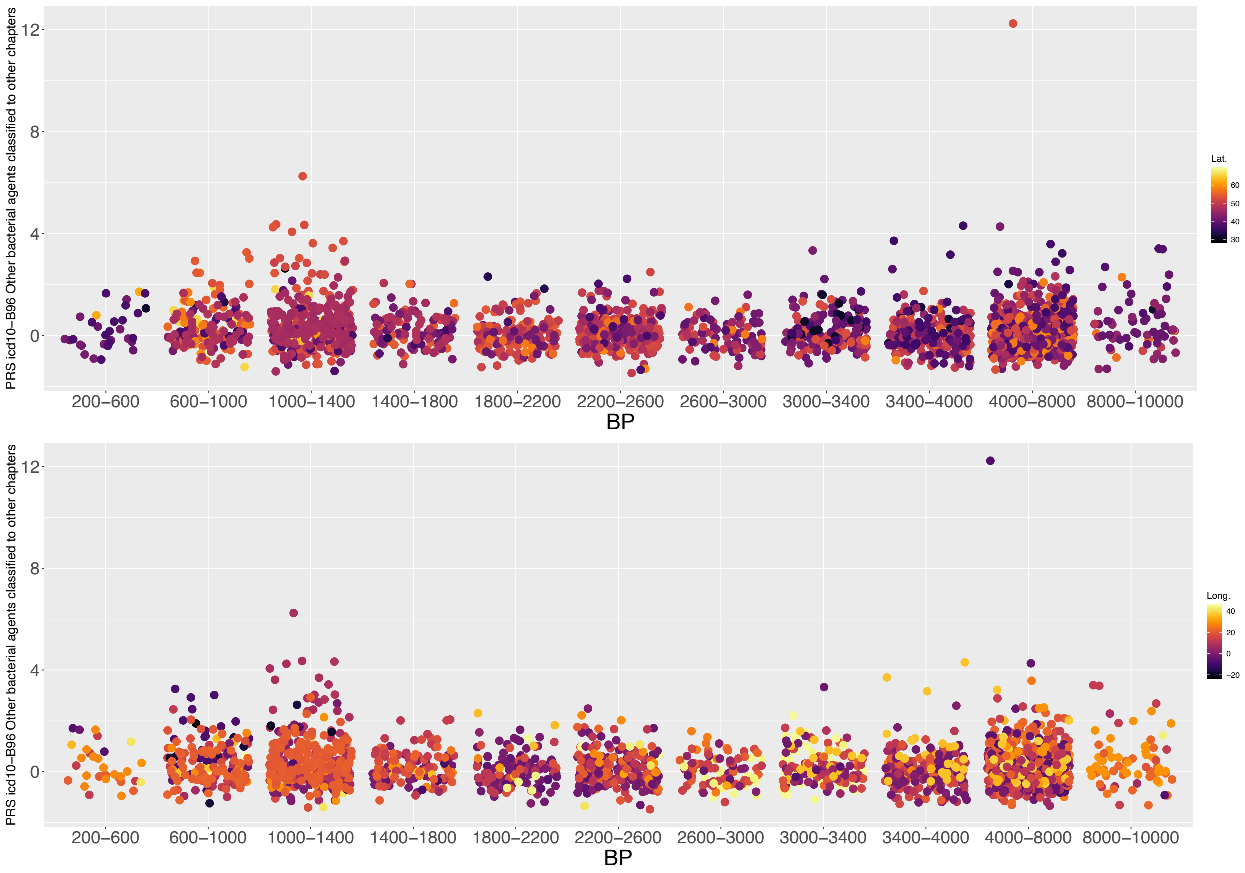


Fig. S4: PRS distribution for *icd10 B96 Other bacterial agents classified to other chapters* binned in 400-year classes. Colors refer to the Latitude (top) and Longitude (bottom) where the individual sample is located according to AADR v.54.1.

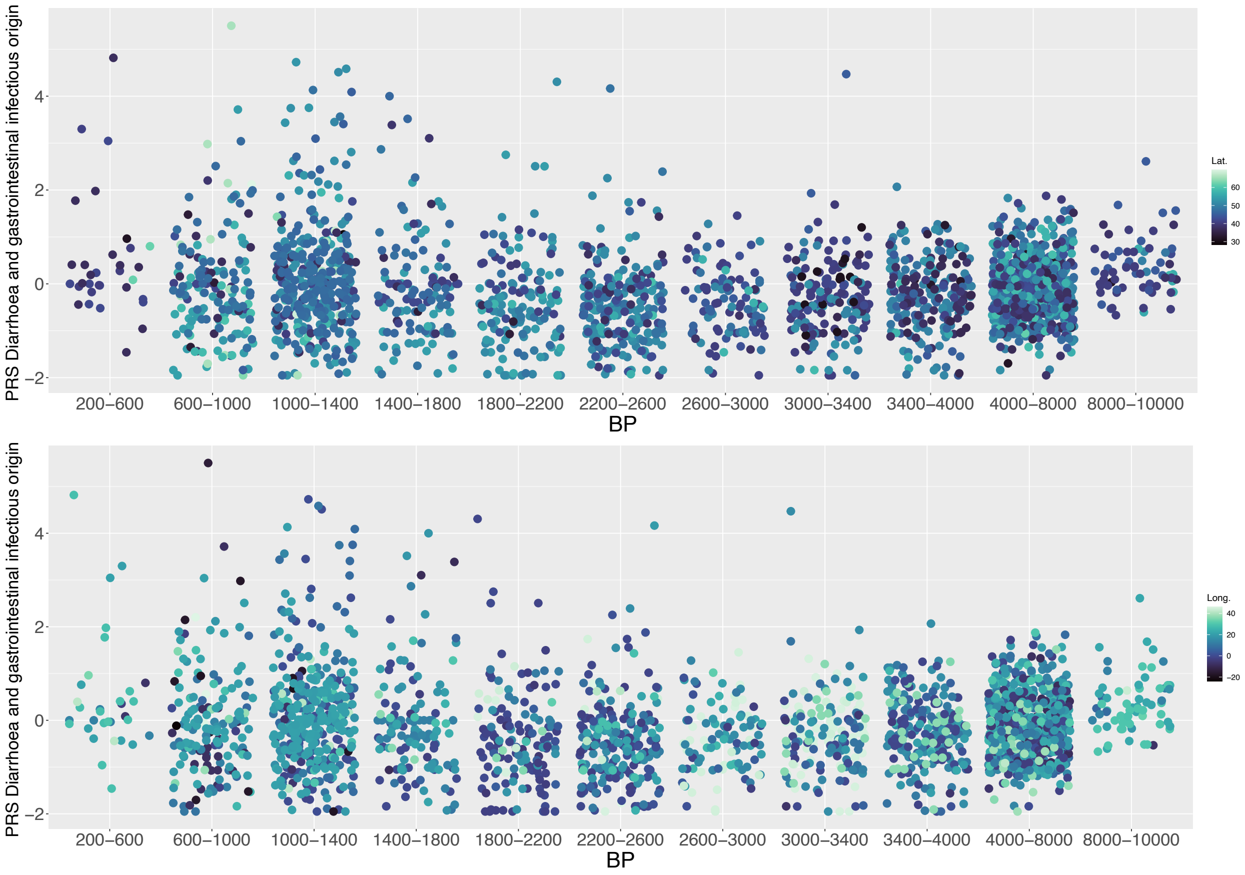


Fig. S5: PRS distribution for *Diarrhoea and gastrointestinal infectious origin* binned in 400-year classes. Colors refer to the Latitude (top) and Longitude (bottom) where the individual sample is located according to AADR v.54.1.

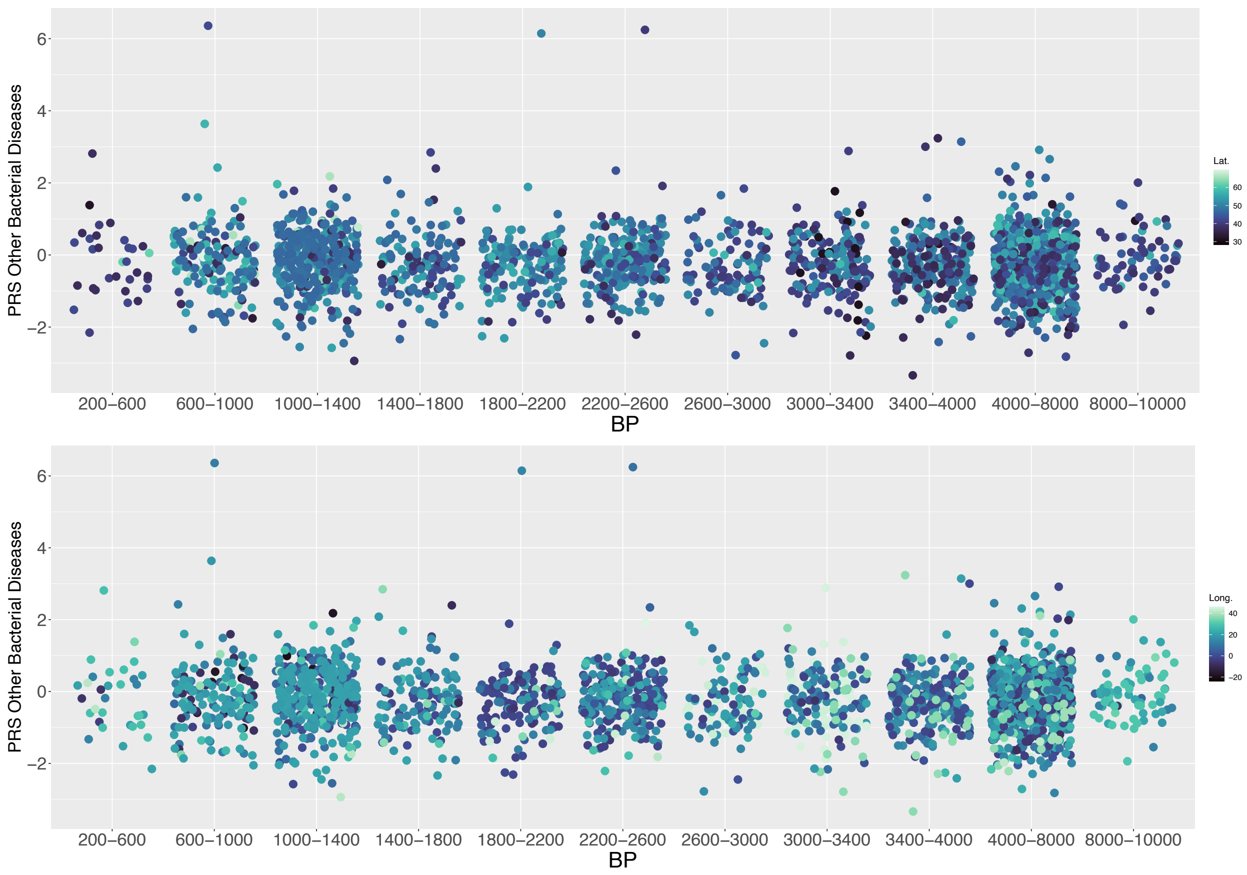


Fig. S6: PRS distribution for *Other Bacterial Diseases* binned in 400-year classes. Colors refer to the Latitude (top) and Longitude (bottom) where the individual sample is located according to AADR v.54.1.

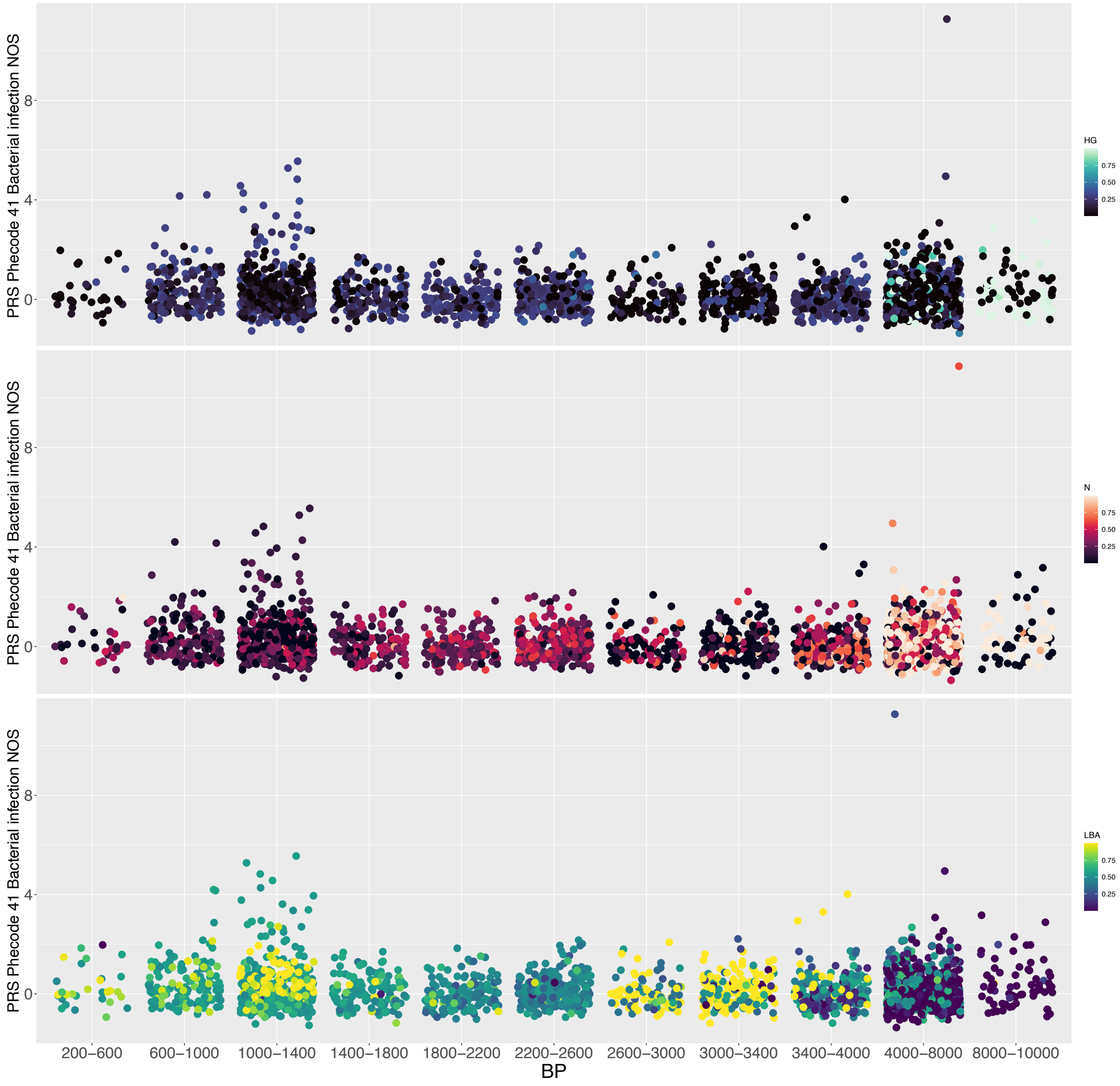


Fig. S7: PRS distribution for *phecode 41 Bacterial infection NOS* binned in 400-year classes. Colors refer to the admixture coefficient with respect to the pre-defined parental classes extrapolated from AADR v.54.1. HG: Hunter-gatherers; N: Neolithic; LBA: Late Bronze Age.

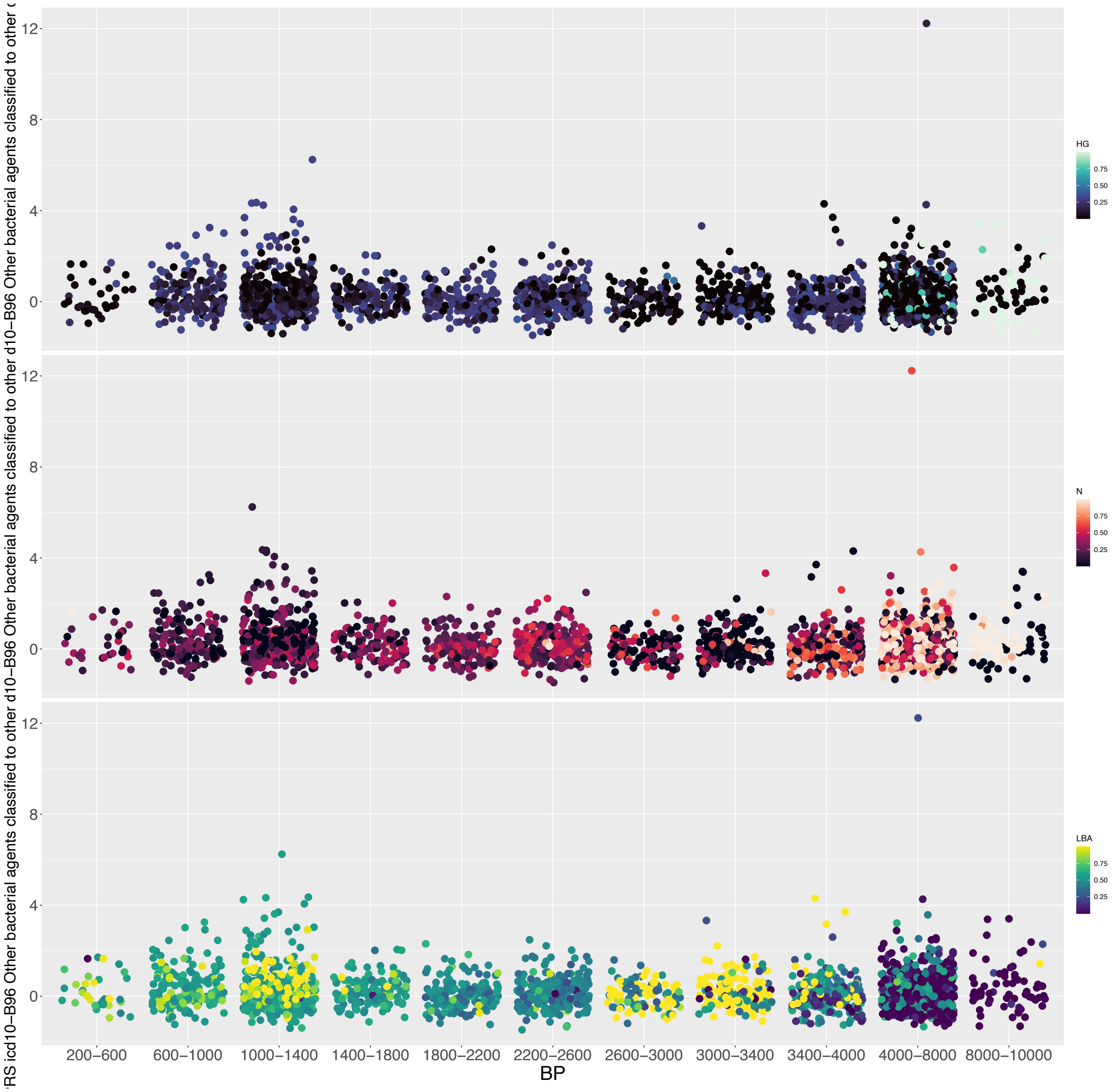


Fig. S8: PRS distribution for *icd10 B96 Other bacterial agents classified to other chapters* binned in 400-year classes. Colors refer to the admixture coefficient with respect to the pre-defined parental classes extrapolated from AADR v.54.1. HG: Hunter-gatherers; N: Neolithic; LBA: Late Bronze Age.

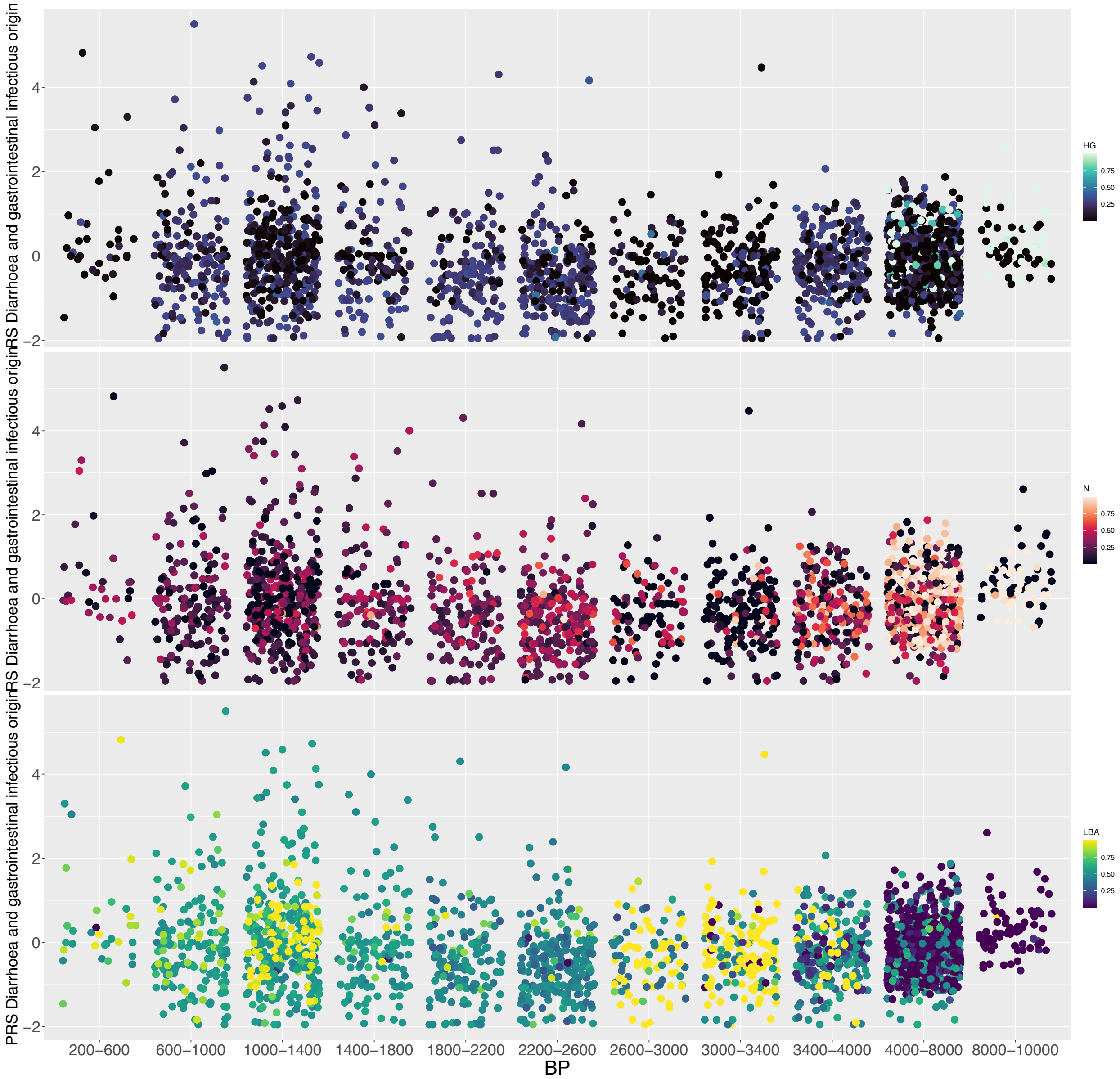


Fig. S9: PRS distribution for *Diarrhoea and gastrointestinal infectious origin* binned in 400-year classes. Colors refer to the admixture coefficient with respect to the pre-defined parental classes extrapolated from AADR v.54.1. HG: Hunter-gatherers; N: Neolithic; LBA: Late Bronze Age.

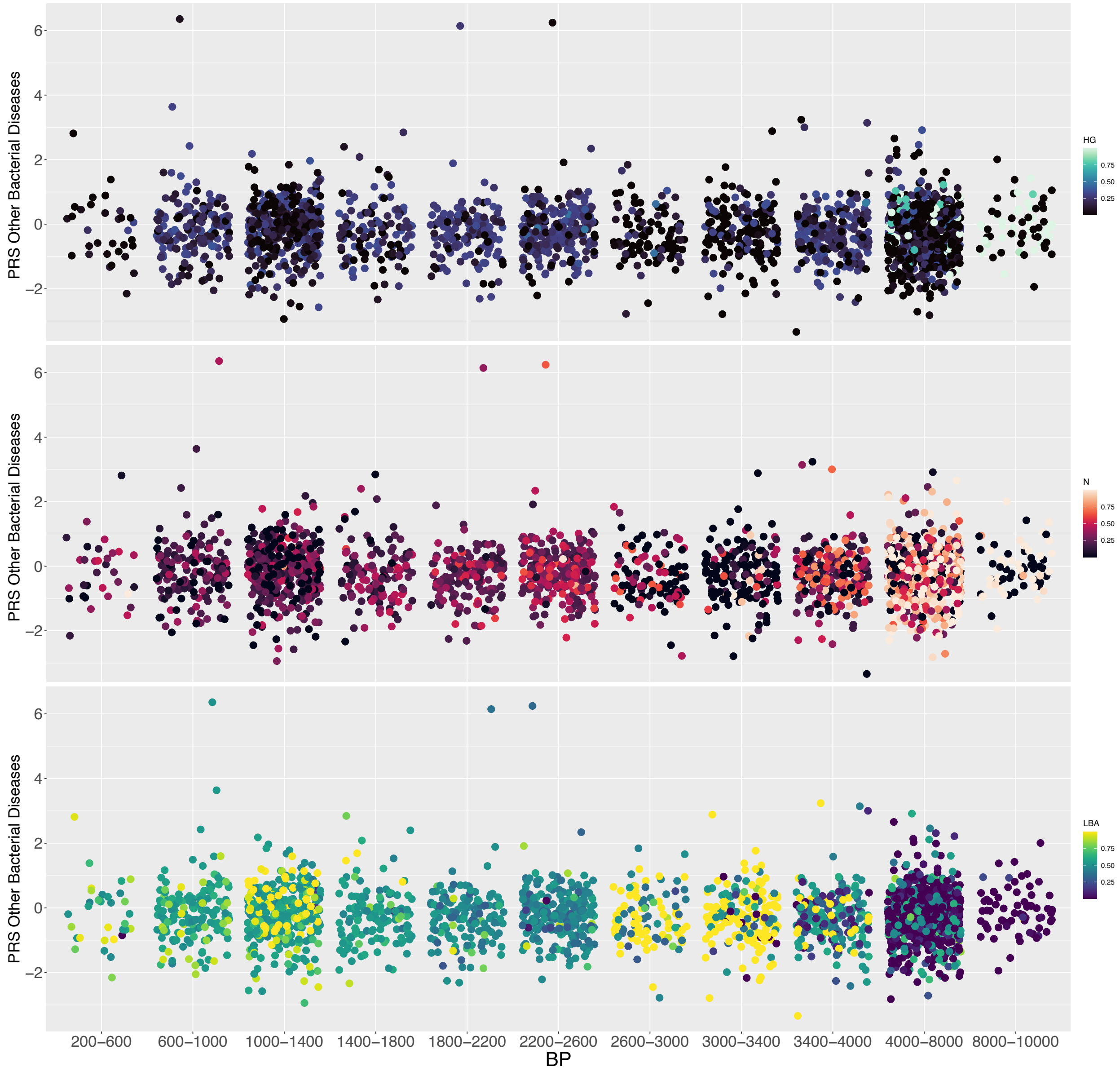


Fig. S10: PRS distribution for *Other Bacterial Diseases* binned in 400-year classes. Colors refer to the admixture coefficient with respect to the pre-defined parental classes extrapolated from AADR v.54.1. HG: Hunter-gatherers; N: Neolithic; LBA: Late Bronze Age.

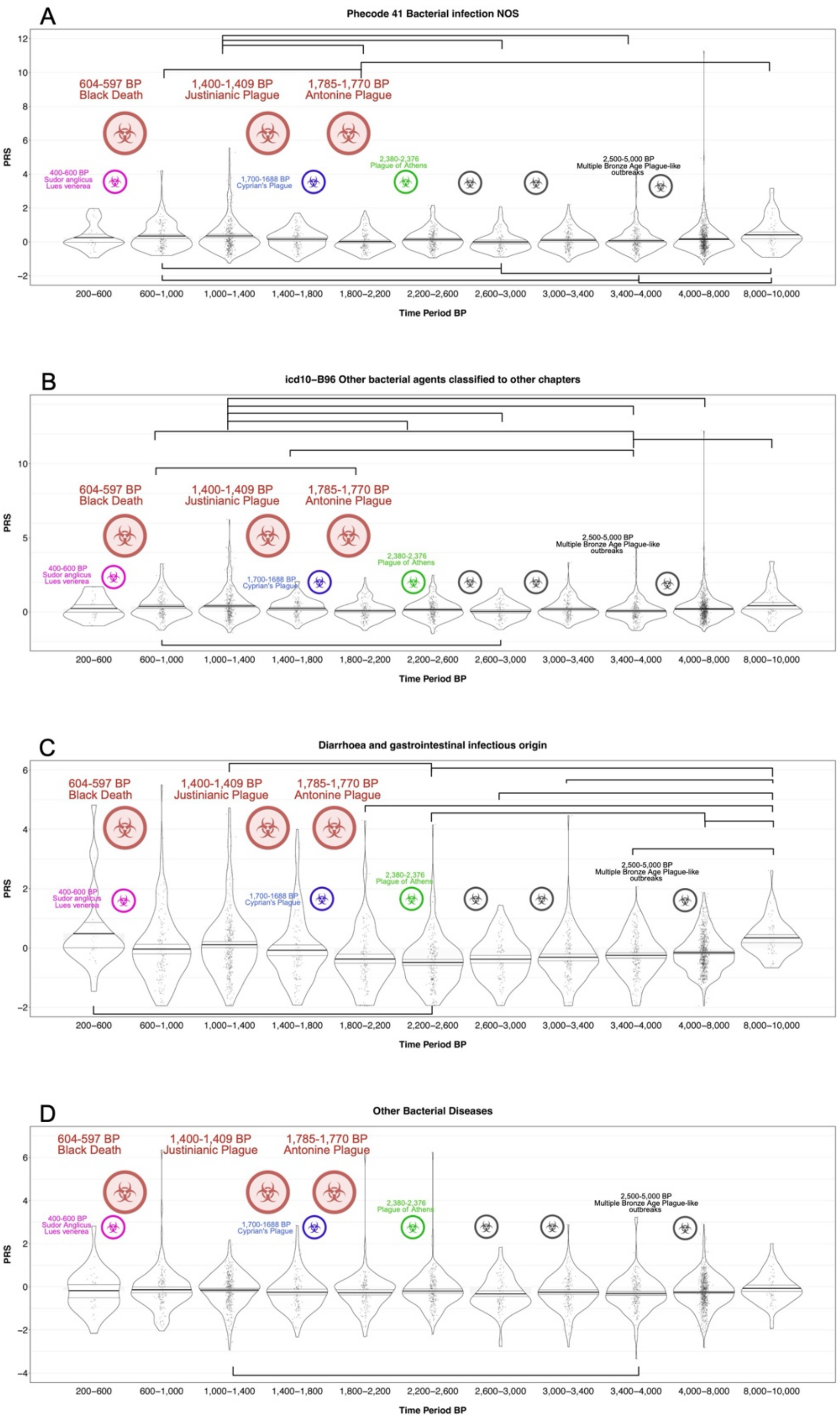


Fig. S11: Distribution of polygenic risk scores (PRS) for four infectious disease-related traits across 400-year time intervals. The X-axis represents the 400-year time intervals in years before present, while the Y-axis shows PRS values. (a) PRS distribution for the UK Biobank (UKB) trait

*“Phecode 41: Bacterial infection, not otherwise specified (NOS)”*. (b) PRS distribution for the UKB trait *“ICD-10 B96: Other bacterial agents classified to other chapters”*. (c) PRS distribution for the FinnGen trait *“Diarrhoea and gastrointestinal infectious origin”*. Only the nine top-ranking significant comparisons are displayed. (d) PRS distribution for the FinnGen trait *“Other Bacterial Diseases”*. The larger biohazard signs refer to the main ID outbreaks discussed in the text or manifestly Plague related outbreaks. The smaller biohazard signs account for other ID outbreaks which cannot be fully detected in our analysis.
